## Supplemental Figures for "Natural variation in salt-induced changes in root:shoot ratio reveals SR3G as a negative regulator of root suberization and salt resilience in Arabidopsis"

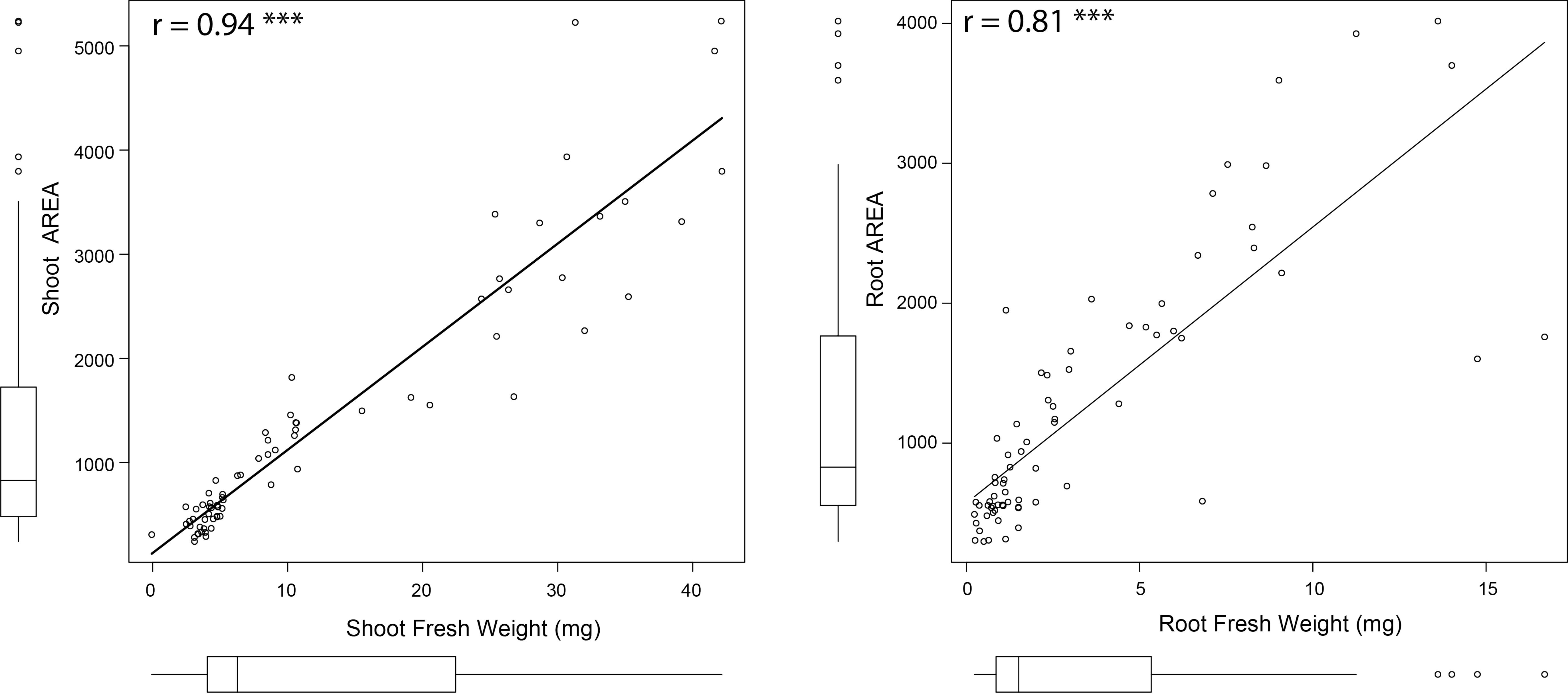


**Figure S1. Evaluation of the tool's precision for estimating seedling size.** The tool used to quantify root and shoot size from the agar plates was tested on the Arabidopsis Col-0 seedlings exposed to various concentrations of salt stress (0, 75 and 125 mM NaCl). The fresh weight of root and shoot of individual plants were recorded at the last day of experiment (8 days after transfer to treatment plates, 12 days after germination), and the correlation analysis between individual organ's fresh weight and area was performed in R. The Correlation co-efficient and the p-value for individual correlations were calculated in R using stats package.

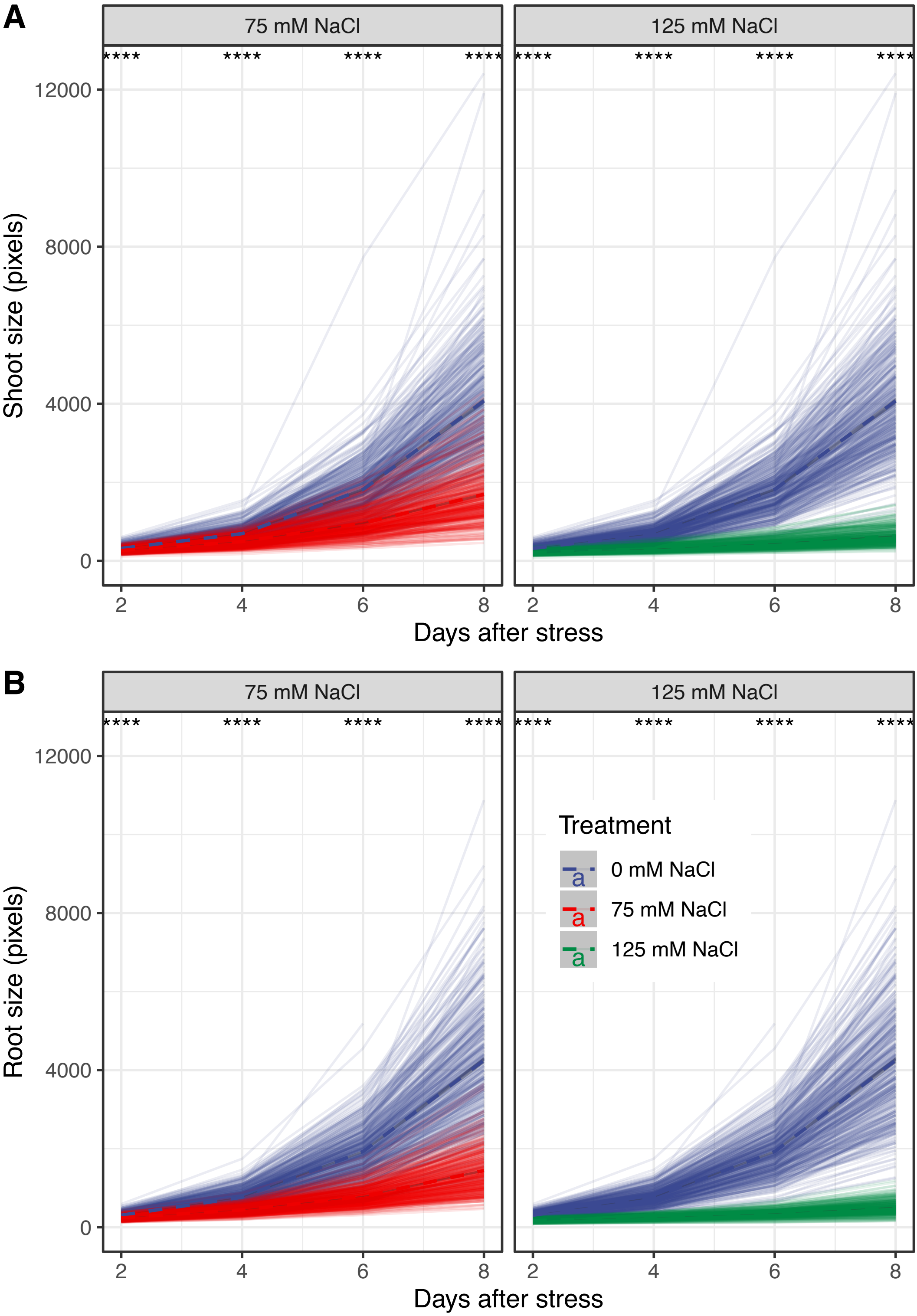


**Figure S2. Salt stress reduced the increase in root and shoot area over time across HapMap accessions.** Four-days-old seedlings exposed to 0, 75 and 125 mM NaCl grown on agar plates were scanned every 2nd day and analyzed for root and shoot area using a custom tool. The change in genotype-specific **(A)** shoot and **(B)** root area was calculated in R using stats package. The thick dashed lines represent population-wide average in root and shoot area over the time of the experiment, while individual transparent lines indicate change in root and shoot area for individual accession. The differences between control and individual salt stress treatments were evaluated using ANOVA. The *, **, *** and **** represent p-values below 0.05, 0.01, 0.001 and 0.0001 respectively.

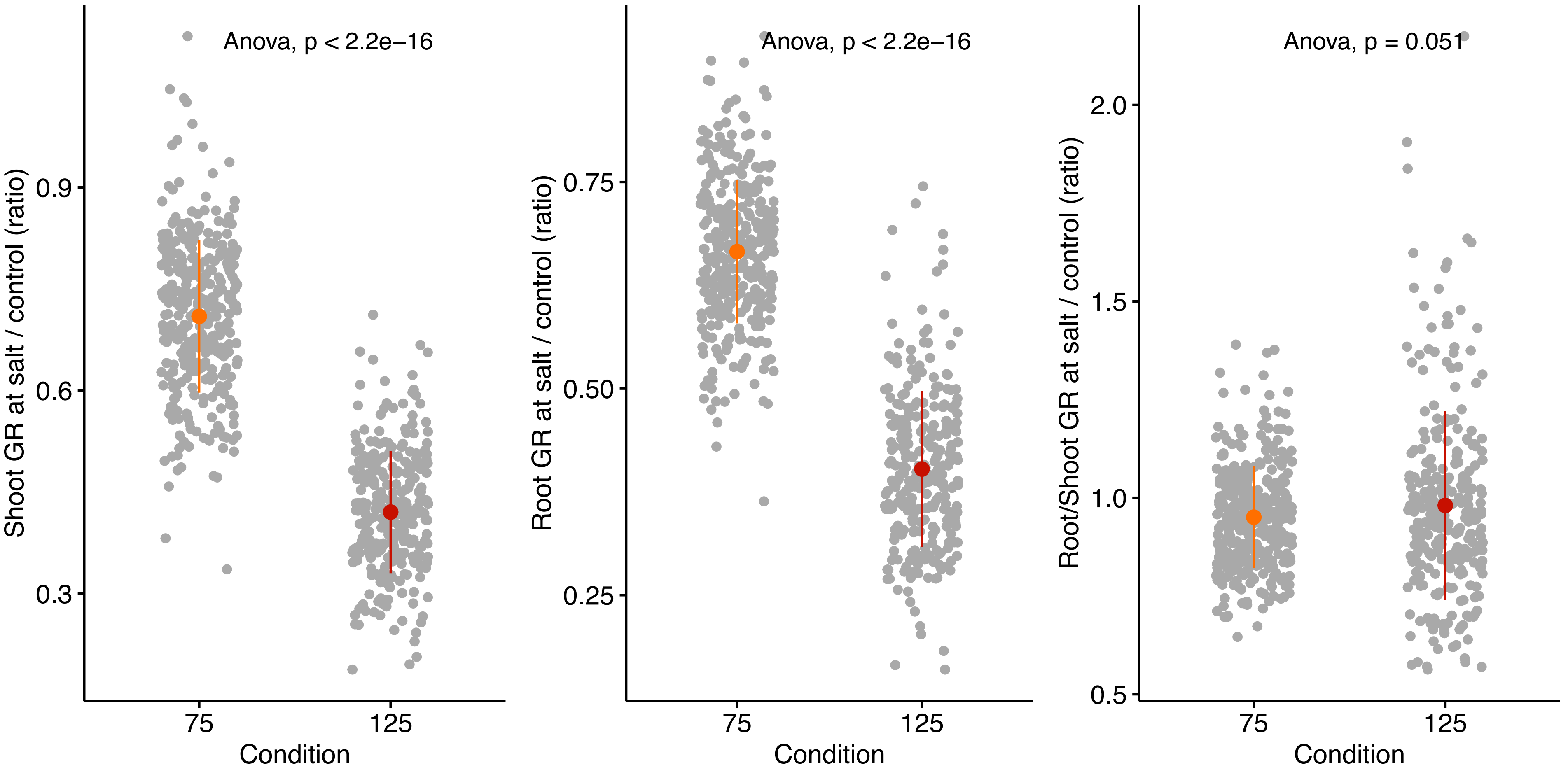


**Figure S3. Salt stress induced changes in relative growth of shoot, root and root:shoot ratio.** The stress tolerance index was calculated for shoot, root and root:shoot ratio by dividing genotype specific value recorded at salt stress over the genotypic mean of the trait recorded under non-stress conditions. Individual points represent individual genotypes. The population mean for 75 and 125 mM NaCl treated plants is represented by the colored point, whereas the population-wide standard error by whiskers.

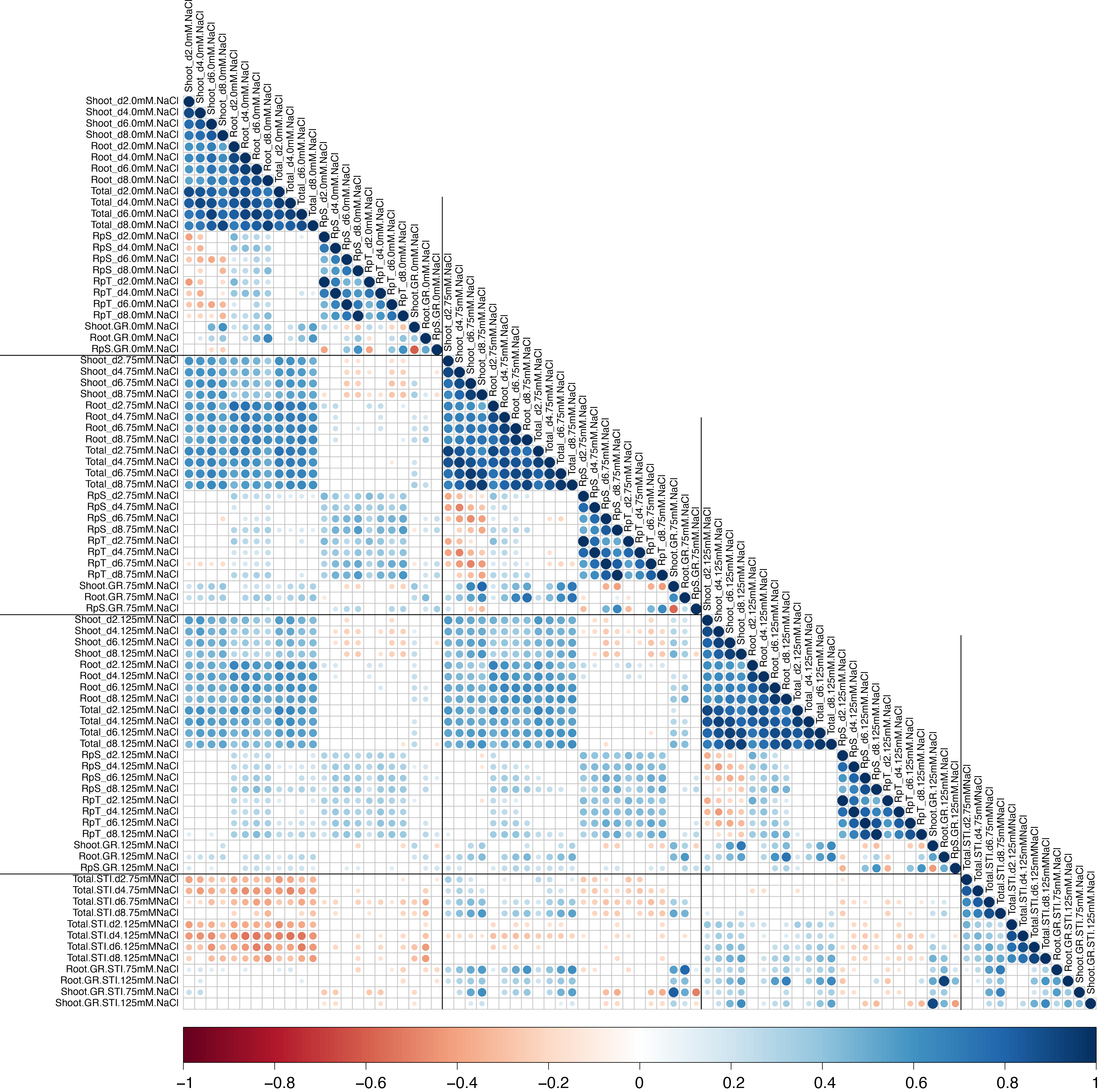


**Figure S4. The correlation between all measured traits and calculated stress tolerance indices (STI).** The correlations between all measured and calculated traits was performed using the Pearson's correlation coefficient. All non-significant correlations were replaced by blank circles. The size and color of individual circles indicates the strength and significance of individual correlation between two traits. Blue and red shades indicate positive and negative correlations respectively.

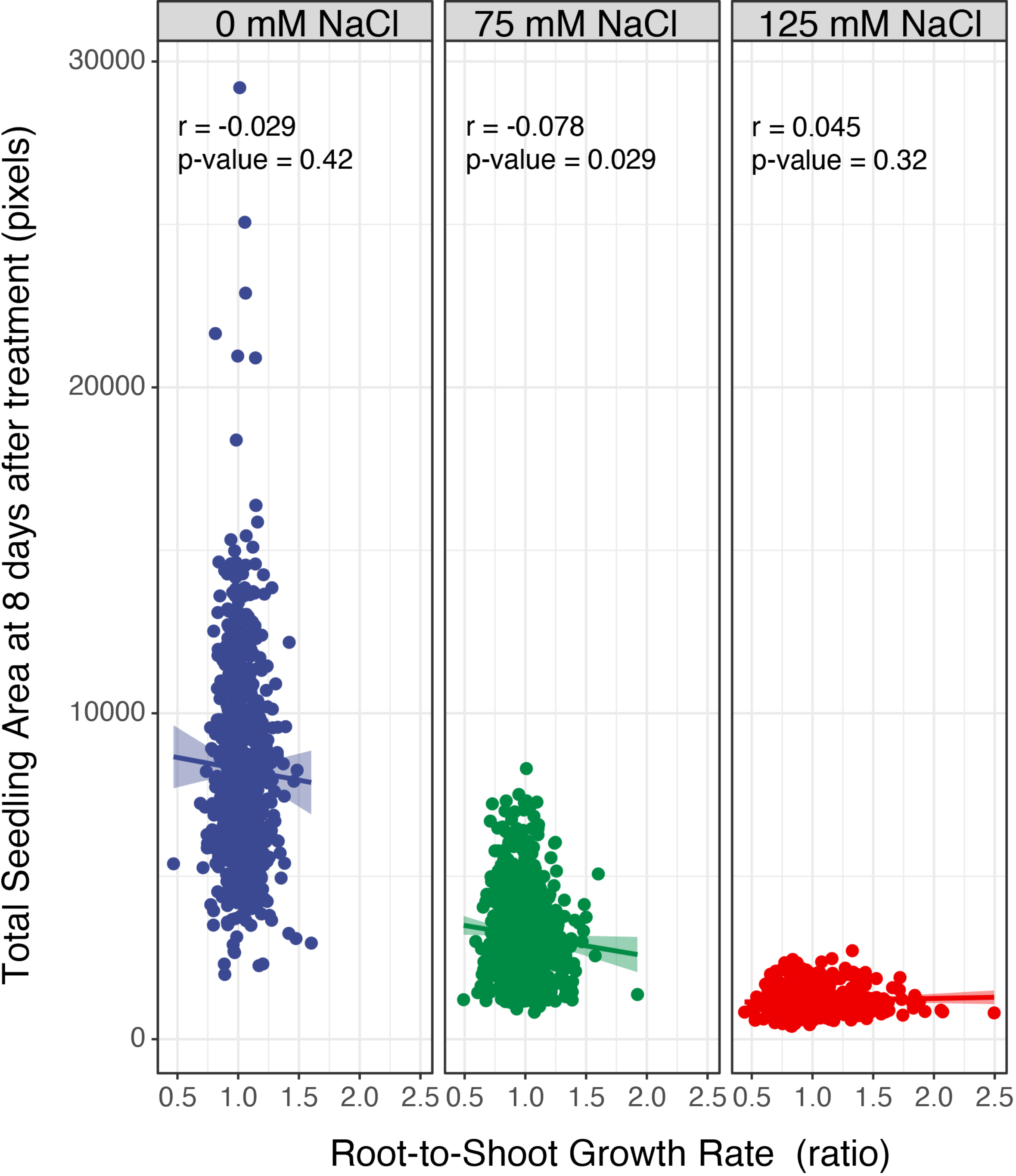


**Figure S5. The correlation between total seedling area and salt-induced changes in the root-to-shoot ratio.** The correlation between the total seedling area quantified using the custom tool at the last day of experiment (8 days after transfer and 12 days after germination), and the ratio of root-and-shoot growth rate was examined across Arabidopsis HapMap population and various concentrations of salt stress (0, 75 and 125 mM NaCl). The correlation analysis was performed in R, and the Correlation co-efficient (r) and the p-values for individual correlations were calculated in R using stats package.

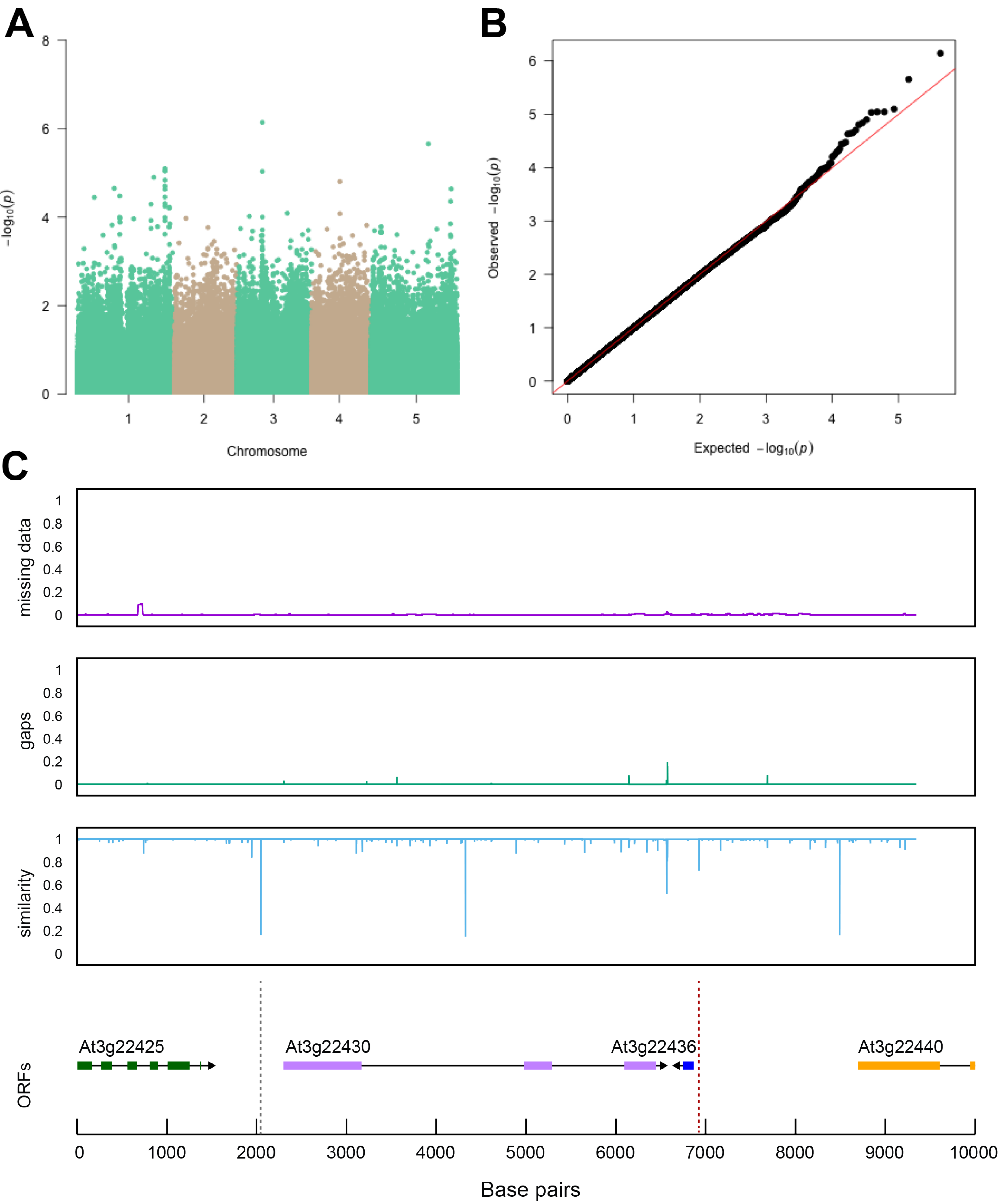


**Figure S6. Region around NTL8 is associated with salt stress-induced changes in root-per-shoot growth.** The salt induced changes in root-to-shoot and shoot-per-total seedling area ratios were used as an input for GWAS. **(A)** Manhattan plot represents the associations of root-per-shoot growth factors recorded at 75 mM NaCl with 250 k SNPs while **(B)** represents the QQ plot for this association study. **(C)** Significant associations were found with the SNPs forming a locus on chromosome 2 in and around AT2G27300, encoding a NAC-domain containing transcription factor (NTL8). The natural variation in the LD region was studied in 162 accessions sequenced by 1001 Genomes Project. The upper panel represents portion of the missing data, upper middle panel represents deletions relative to Col-0, while lower middle panel represents the sequence similarity compared to Col-0. The bottom panel represents the open reading frames (ORFs) within the LD, and the location of associated SNPs is indicated with the dashed lines. Red dashed lines represent associations above Bonferroni threshold in 250k SNP mapping, while grey lines represent associations with -log10(p-value) > 5 in the 4M SNP mapping.

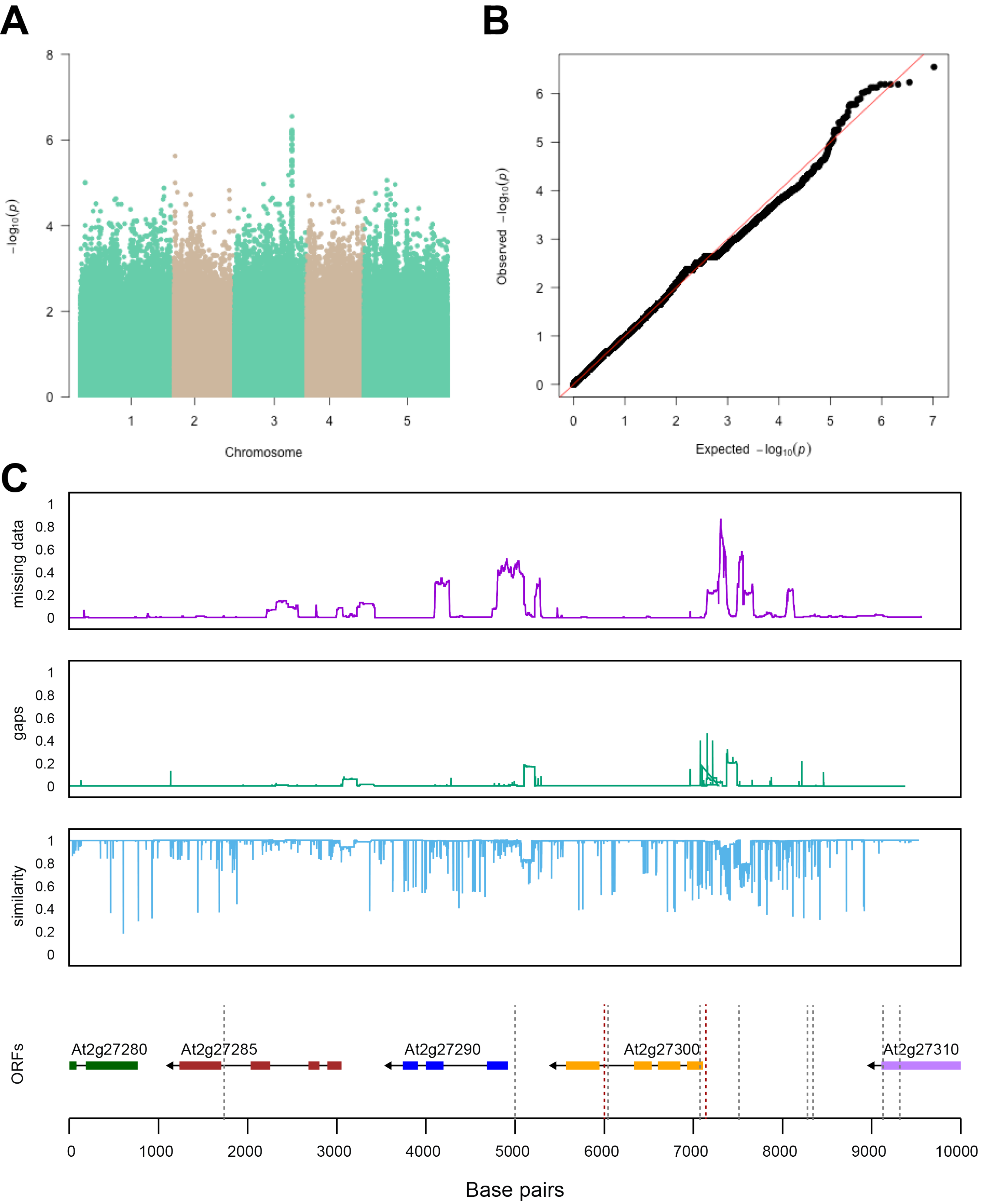


**Figure S7. Region around unknown gene is associated with salt stress-induced changes in relative changes in shoot-per-total seedling area.** The salt induced changes in root-to-shoot and shoot-per-total seedling area ratios were used as an input for GWAS. **(A)** Manhattan plot represents the associations of relative change (125 mM / 0 mM) in shoot-per-total seedling area at 4 days after transfer to 125 mM NaCl, while **(B)** represents the QQ plot for this association study. **(C)** Significant associations were found with the SNPs forming a locus on chromosome 3 in and around AT3G22430, encoding an unknown gene encoding a domain of unknown function. The natural variation in the LD region was studied in 162 accessions sequenced by 1001 Genomes Project. The upper panel represents portion of the missing data, upper middle panel represents deletions relative to Col-0, while lower middle panel represents the sequence similarity compared to Col-0. The bottom panel represents the open reading frames (ORFs) within the LD, and the location of associated SNPs is indicated with the dashed lines. Red dashed lines represent associations above Bonferroni threshold in 250k SNP mapping, while grey lines represent associations with -log10(p-value) > 5 in the 4M SNP mapping.

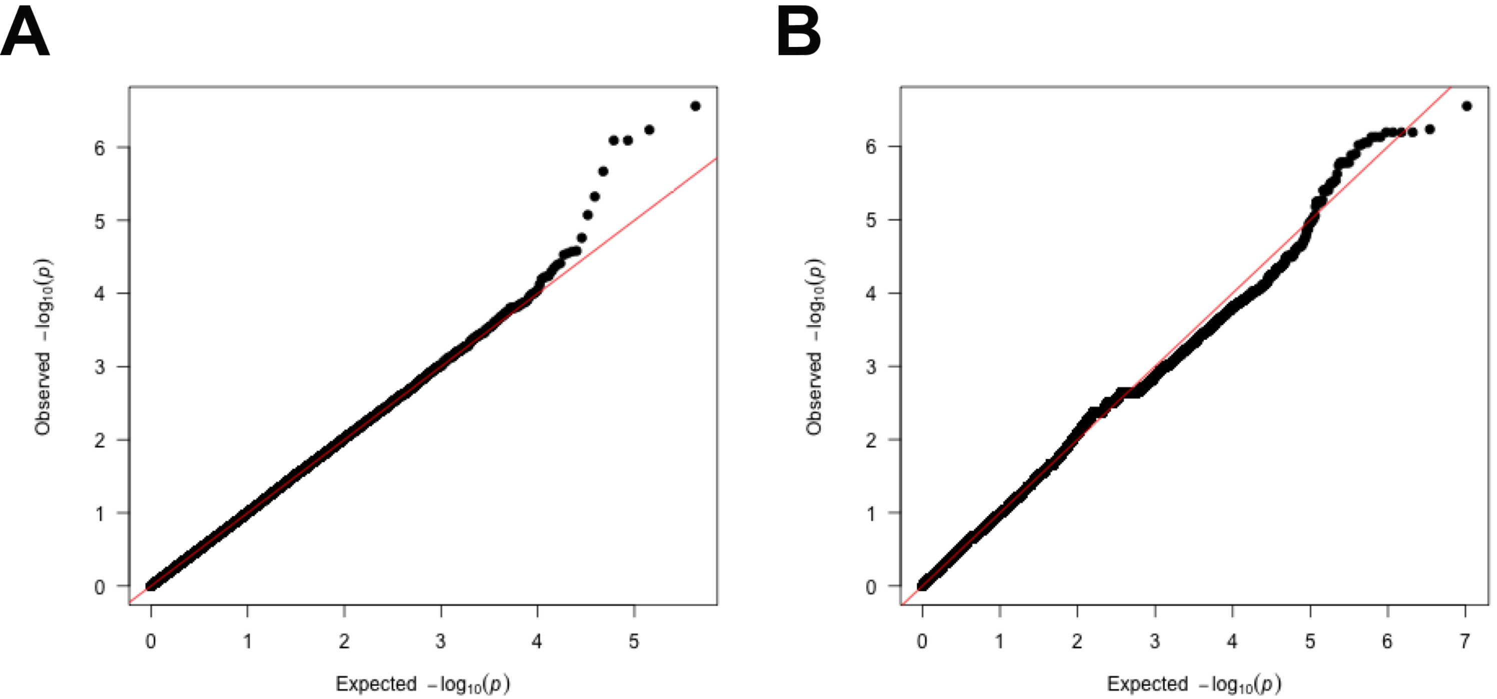


**Figure S8. The QQ-plots for GWAS models with shoot-per-total seedling area.** The QQ-plots for association between shoot-per-total seedling area recorded 6 days after transfer to 125 mM NaCl and **(A)** 250 k SNPs and **(B)** 4 M SNPs. The red line indicates the expected vs. observed ratio of association, while black points indicate the observed associations.

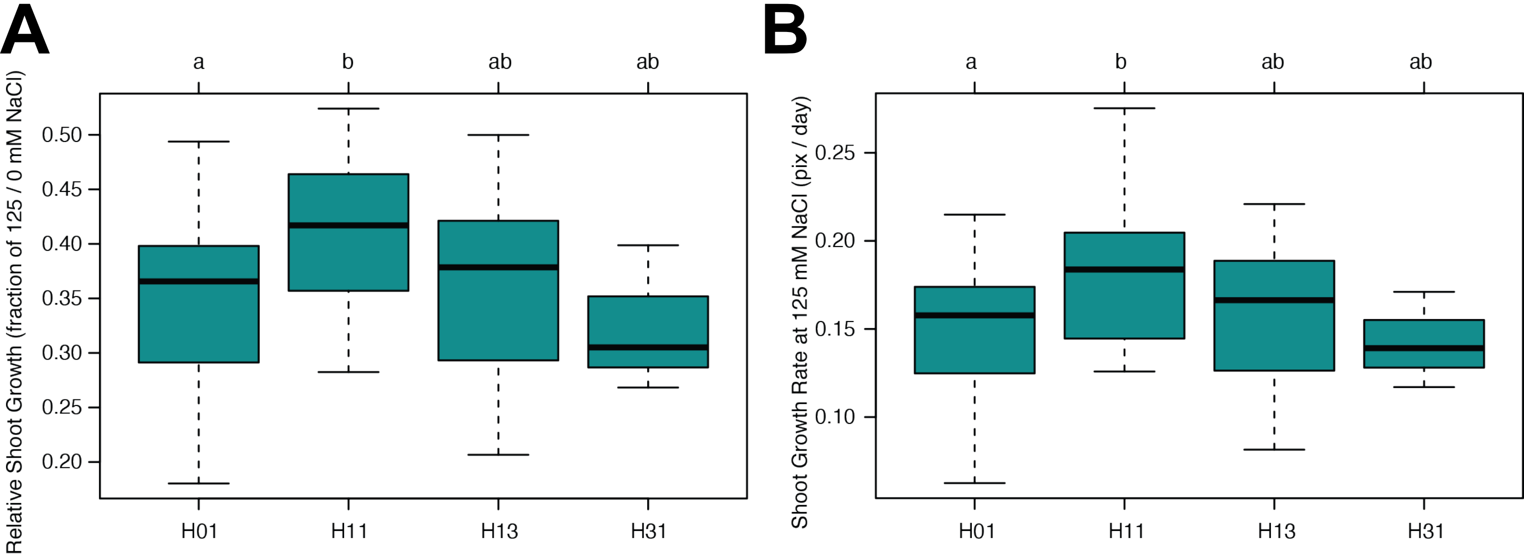


**Figure S9. Haplotype analysis of AT3G50160 (SR3G).** The haplotype analysis performed using SNPs located within the coding region of the SR3G (AT3G50160) revealed significant differences in **(A)** Relative shoot growth (as fraction of shoot growth at 125 mM NaCl and 0 mM NaCl) and **(B)** Shoot growth at 125 mM NaCl. The significant differences between individual haplotype groups (each haplotype represented by at least 3 accessions), were tested using ANOVA with Tukey HSD to identify significantly different groups. Significant differences were identified between Haplotype groups 1 (represented by 46 accessions, including Col-0) and 11 (represented by 11 accessions, including Blh-1).

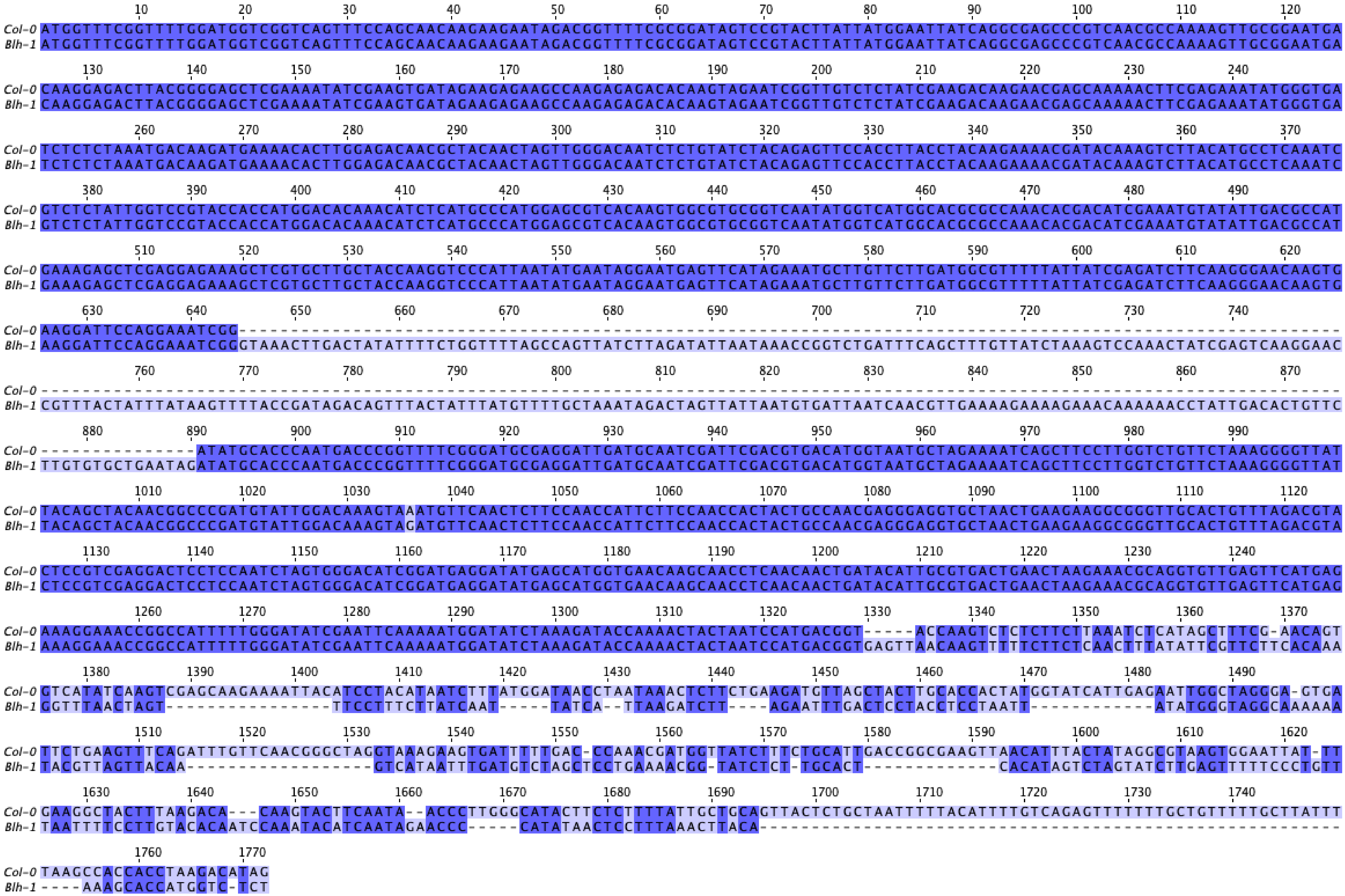


**Figure S10. Nucleotide sequence alignment of SR3G CDS between Col-0 and Blh-1 alleles.** The nucleotide sequences from *SR3G* CDS were obtained from Col-0 and Blh-1 cDNA, and compared to Col-0 cDNA sequence on TAIR. The cDNA sequences were subsequently aligned in JalView using ProbconsWS alignment. The intensity of purple hue indicates the nucleotide identity, while dash marks indicate insertion in Blh-1 compared to Col-0 allele.

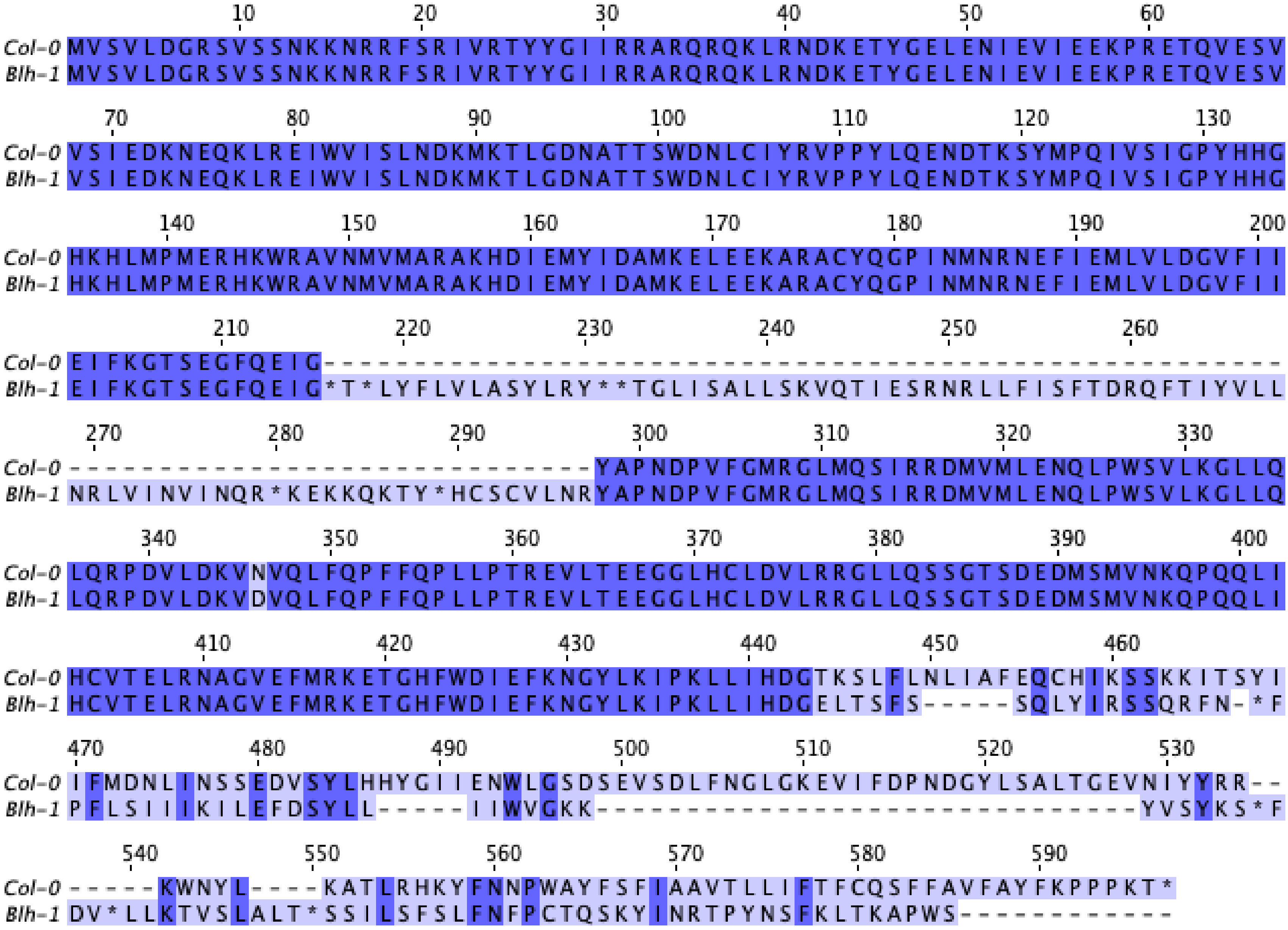


**Figure S11. Amino acid sequence alignment of SR3G CDS between Col-0 and Blh-1 alleles.** The amino acid sequences from *SR3G* CDS were obtained by translating Col-0 and Blh-1 cDNA into amino acid sequence in JalView using Standard settings. The amino acid sequences were subsequently aligned in JalView using ProbconsWS alignment. The sequence length markers for are indicated above the aligned sequence. The intensity of purple hue is indicating the nucleotide identity, while dash marks indicate insertion in one of the alleles. * indicate STOP codons.

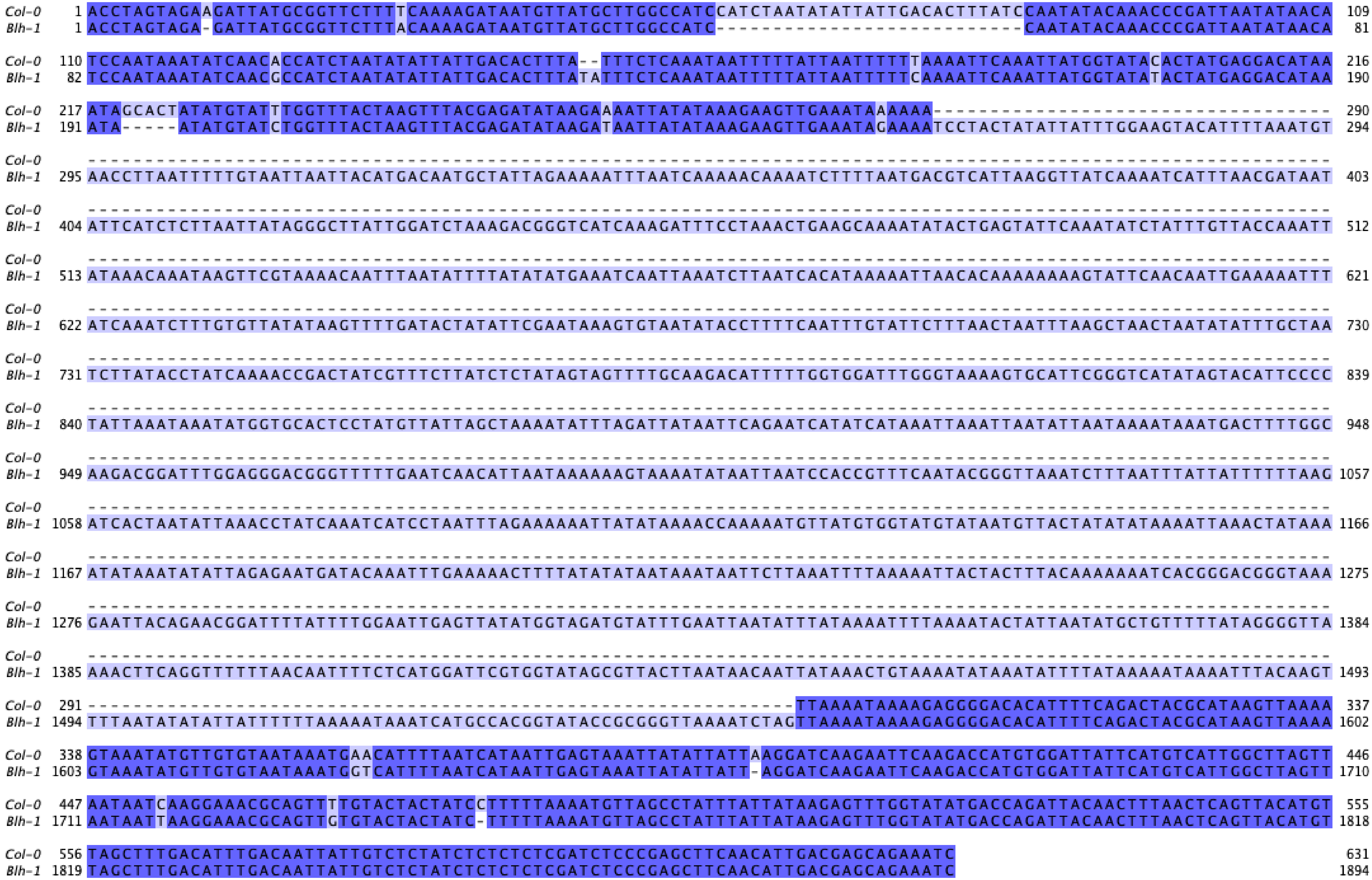

**Figure S12. Sequence alignment between Col-0 and Blh-1 alleles for *SR3G* promotor.** The nucleotide sequences from *SR3G* promotor were obtained from Col-0 and Blh-1 genomic DNA, and compared to Col-0 sequence on TAIR. The gDNA sequences were subsequently aligned in JalView using ProbconsWS alignment. The intensity of purple hue is indicating the nucleotide identity, while dash marks indicate insertions.

A

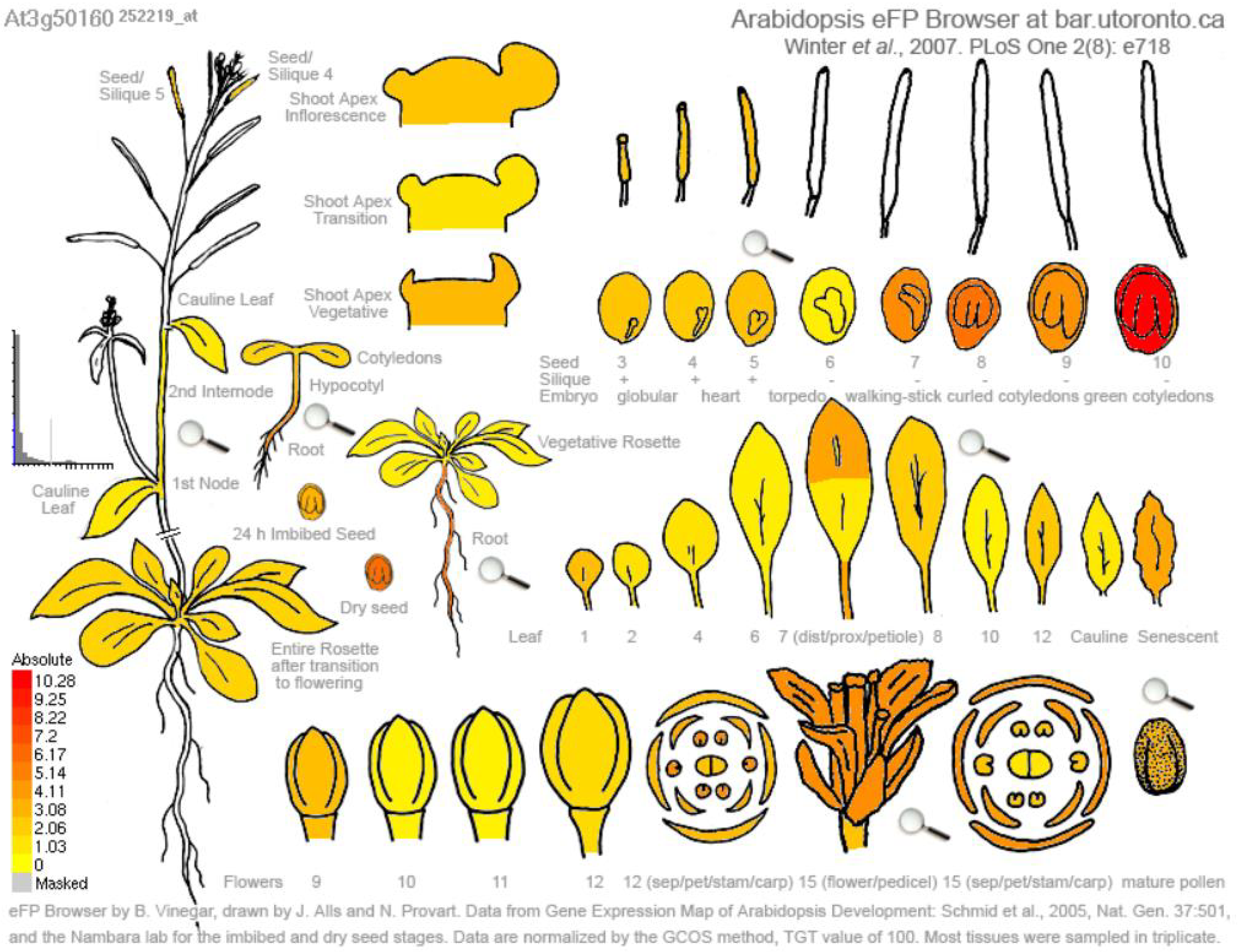


B

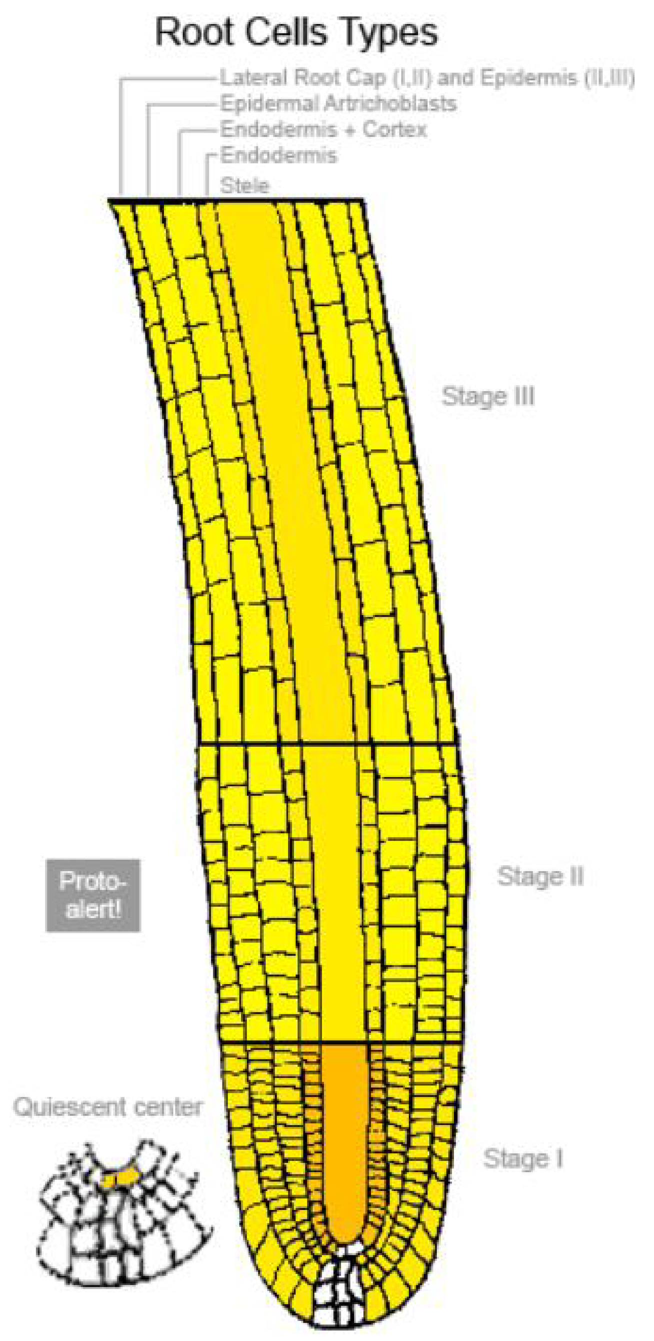


C

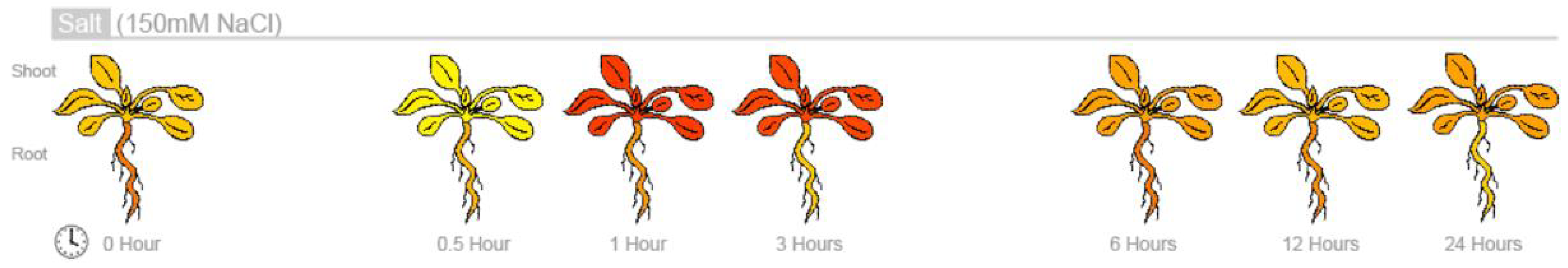


**Figure S13. Gene expression profile of *SR3G* across various developmental stages and salt stress.** The absolute gene expression of SR3G is depicted **(A)** across various developmental stages, **(B)** root cell types, and **(C)** in response to salt stress. The images were acquired through the eFP Browser using data sources specified for the developmental map, abiotic stress, and tissue specific.

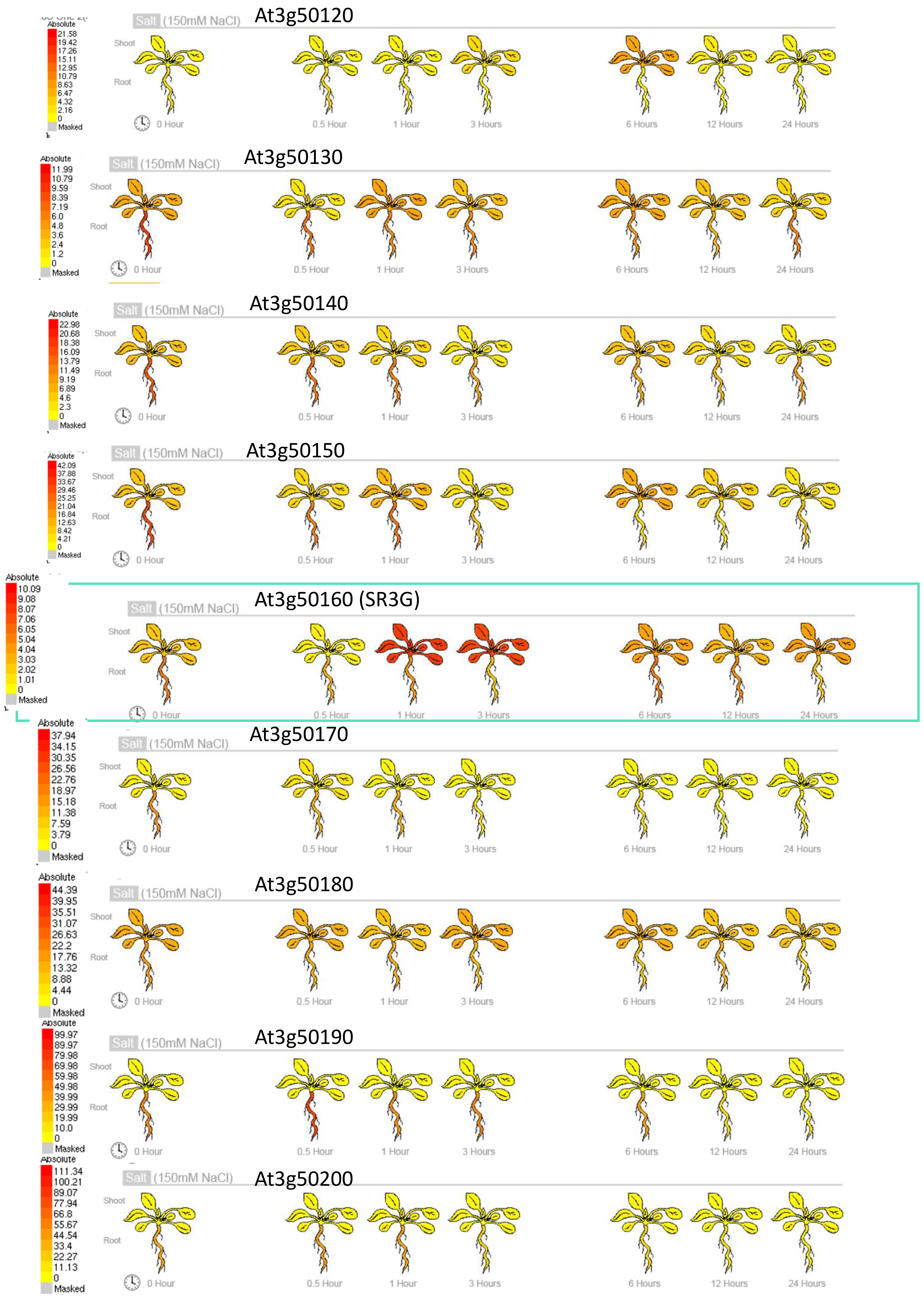


**Figure S14. Gene expression profile of other DUF247s under salt stress.** Absolute gene expression of other DUF247s is illustrated in response to salt-induced changes. The images were acquired through the eFP Browser using data sources specified for the abiotic stress. The *SR3G* is shown in a green box for comparison.

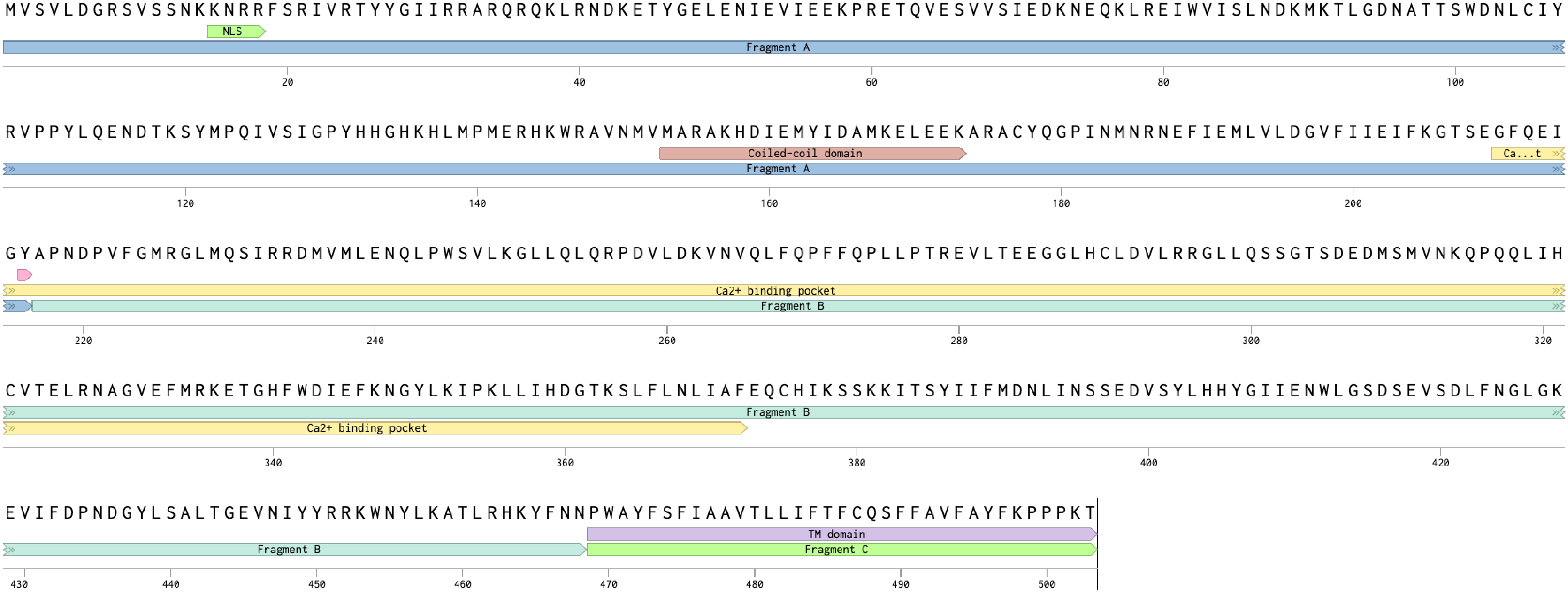


**Figure S15. *SR3G* predicted protein domains.** The *SR3G* protein sequence (Col-0 allele) was used to identify predicted protein domains. The identified domains include: nuclear localization signal (NLS): K15-F19; coiled-coil domain: M153-K173; potential $Ca^{2+}$ binding pocket: G210-F372; and transmembrane domain P469-T503. Informed by these predicted protein domains, we designed SR3G Fragment A, B and C for subsequent localization studies and functional characterization. The pink mark at Y216 indicates a position of a premature STOP codon in Blh-1 allele.

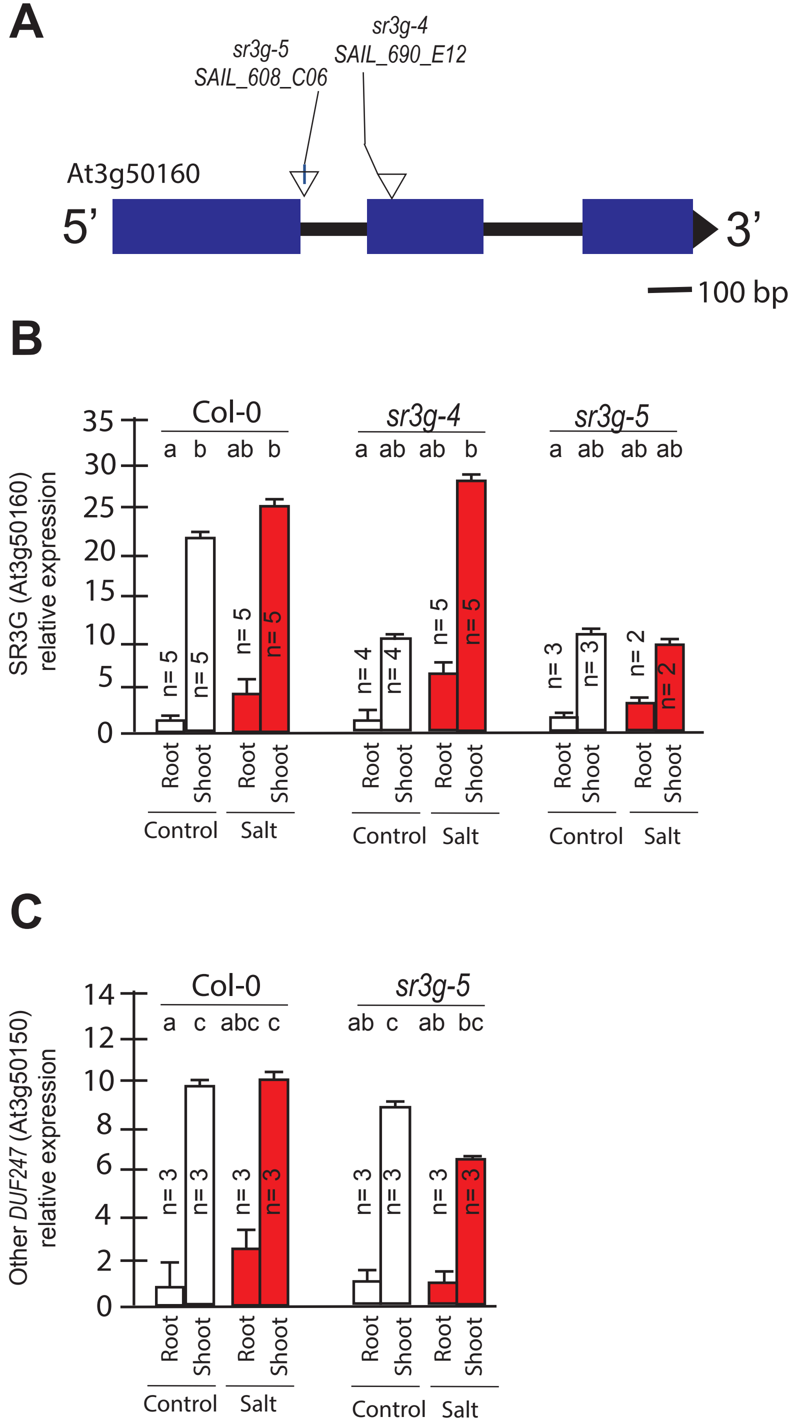


**Figure S16. The expression of DUF247-150 remains unaltered in the *sr3g* mutant. (A)** Schematic diagram of main *SR3G* gene model and the location of T-DNA insertions. The location of T-DNA insertions is shown for *sr3g-4*, and *-5* mutants using triangles. **(B)** RT-qPCR showing expression of *SR3G* gene (AT3G50160) and **(C)** its neighboring gene, DUF247-150 (At3g50150) in Col-0 and two *sr3g* mutants. RT-qPCR analyses were conducted using seedlings grown on 1/2x MS for 4 days and then followed by transferring to the treatment plates with or without 75 mM NaCl for one week. Mean values are shown ± SE, with the number of replicates (n) shown in each graph. AT4G04120 was used as reference gene for normalization. Significance was determined by the Tukey–Kramer HSD test in JMP. Levels not connected by same letter are significantly different.

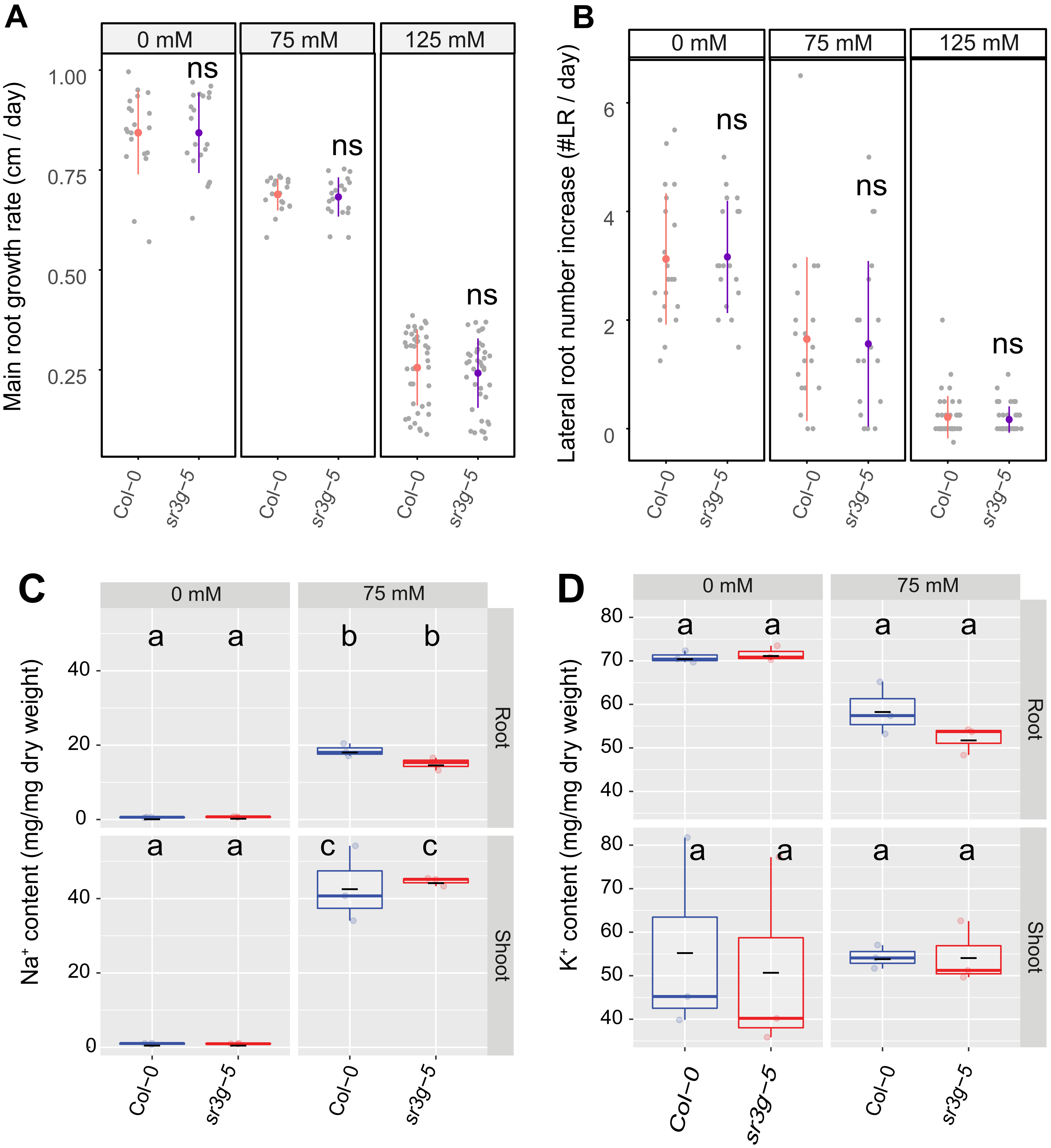


**Figure S17.** ***sr3g-5*** **mutant displays no difference in main root length and lateral root numbers compared to Col-0.** Root system architecture analysis of Col-0 and *sr3g-5* plants under various concentrations of NaCl are shown here for **(A)** main root length and **(B)** increase in lateral root number. 13-day-old Col-0 and sr3g-5 genotypes experienced 9 days of salt treatment at indicated concentrations were used here. Each dot in (A) and (B) represents individual replicate per genotype. Statistical analysis was performed between Col-0 and the *sr3g-5* mutant using the Student 's t-test: $^*P < 0.05$. ns stands for not significant. **(C)** $Na^+$ and **(D)** $K^+$ contents in root and shoot of Col-0 and *sr3g-5* after two weeks on 75 mM salt are shown. Statistical analysis in (C-D) was done by comparison of the means for all pairs using Tukey–Krvqamer HSD test. Levels not connected by the same letter are significantly different ($P < 0.05$).

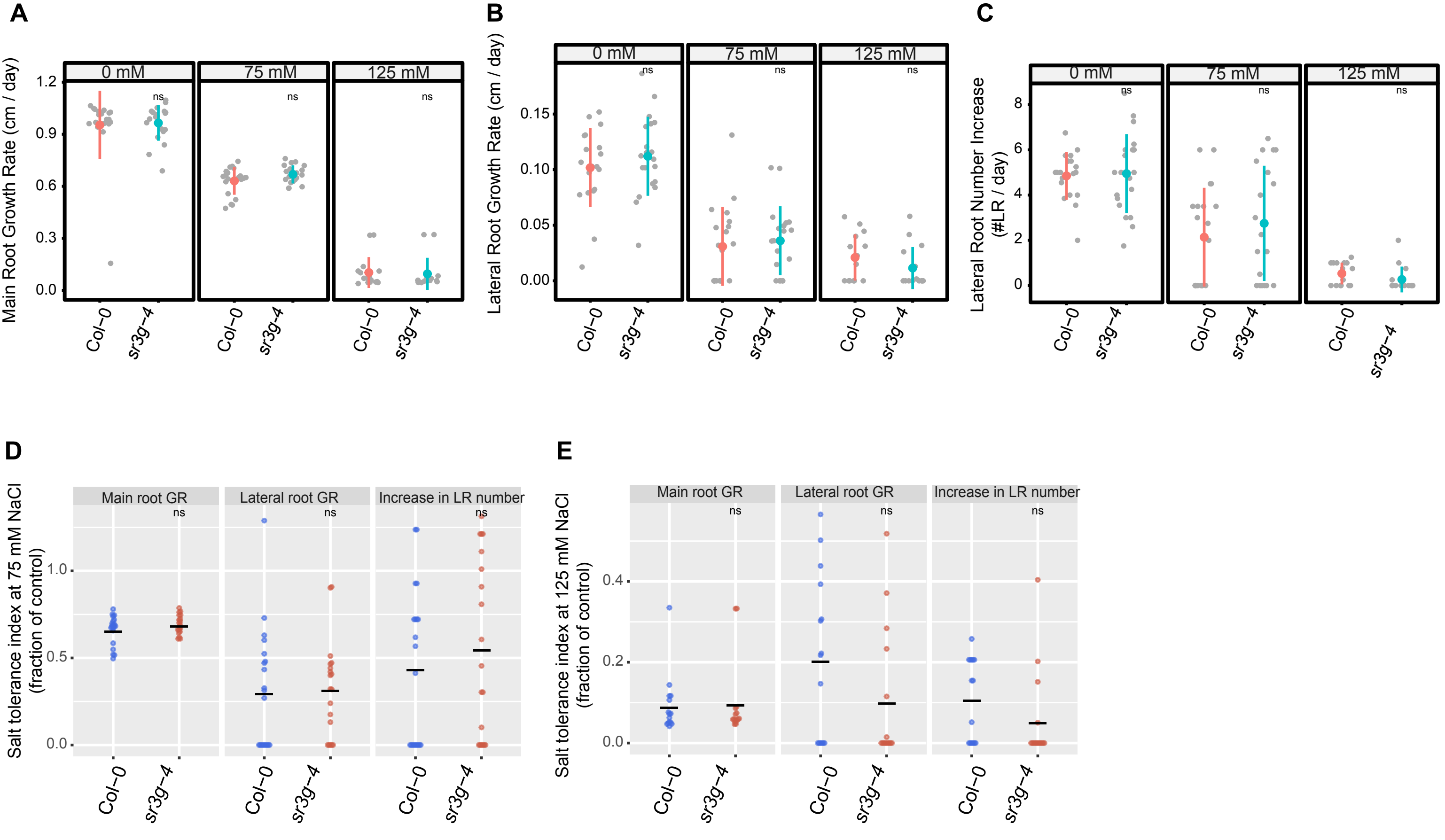


**Figure S18. *sr3g-4* mutant has no alteration in root system architectures compared to Col-0.** Root system architecture analysis of Col-0 and *sr3g-4* plants under various concentrations of NaCl are shown here for **(A)** main root length, **(B)** lateral root length, and **(C)** lateral root number. Salt Tolerance Index (STI) for the main root length, average lateral root length, and lateral root number are shown at **(D)** 75 and **(E)** 125 mM NaCl, respectively. The STI was calculated by dividing the growth rate measured under salt stress by the growth rate measured under control condition for each genotype. Statistical analysis was performed between Col-0 and the *sr3g-4* mutant using the Student ‘s t-test: *P < 0.05.

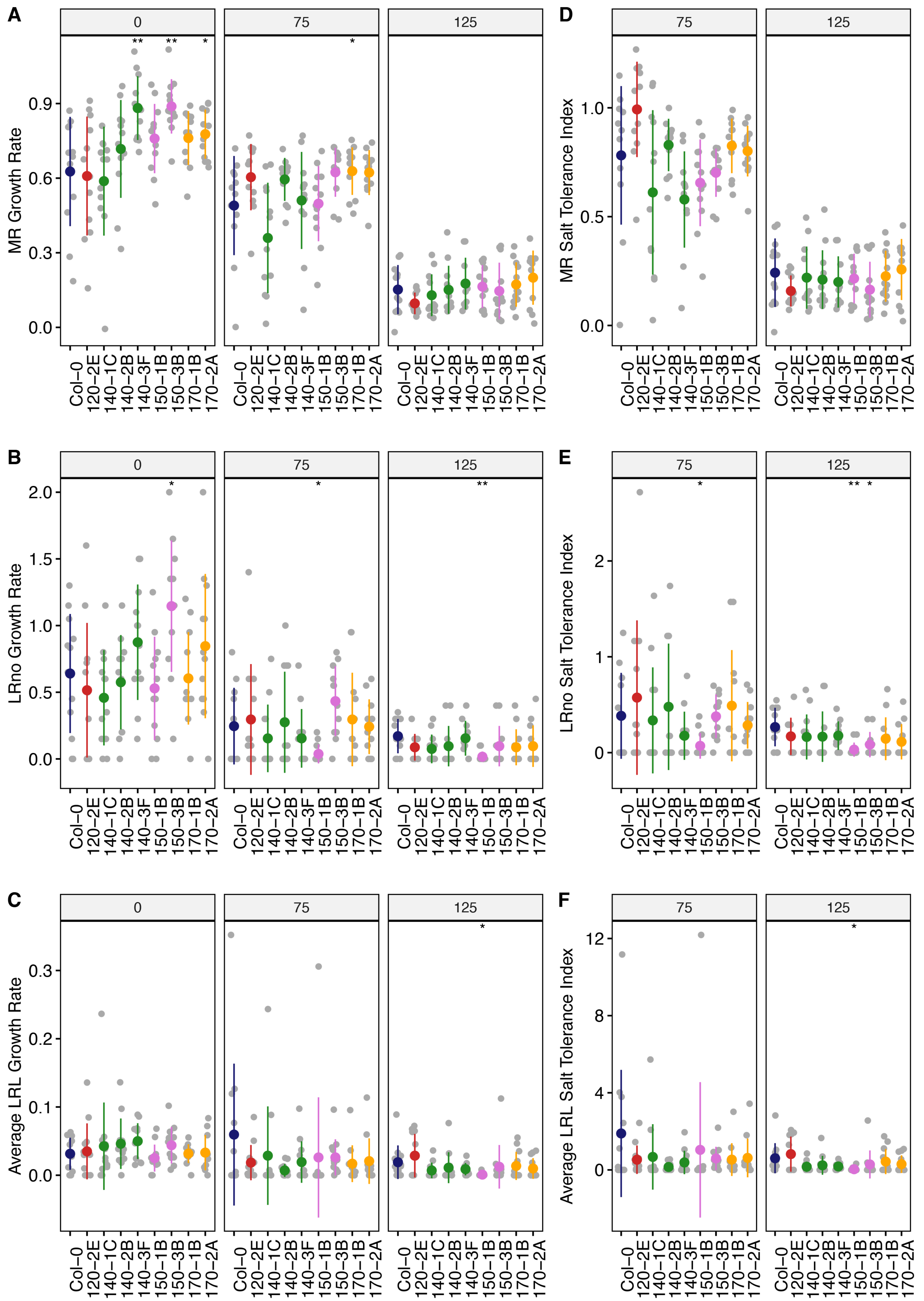


**Figure S19. Root system architecture analysis of neighboring DUF247 mutants under the salt. (A)** Main root length, **(B)** lateral root number, and **(C)** lateral root length are shown for neighboring DUF247s of the *SR3G*, including At3g50120 (shown as 120, with one T-DNA representative, 120-2(SAIL_382_A09)), At3g50140 (shown as 140, with three T-DNA representatives, 140-1 (SALK_044685), 140-2 (SALK_005466), and 140-3 (SALK_122700)), At3g50150 (shown as 150, with two T-DNA representatives, 150-1 (SALK_003824) and 150-3 (SALK_071080)), and At3g50170 (shown as 170, with two T-DNA representatives, 170-1 (SALK_009186) and 170-2 (SALK_112602)) compared to Col-0 under various concentrations of NaCl as indicated. The corresponding Salt Tolerance Index (STI) was calculated for **(D)** main root growth, **(E)** increase in lateral root number, and **(F)** lateral root growth rate. Statistical analysis was performed between Col-0 and the other mutants using the Student 's t-test: *P < 0.05. The root system architecture analysis was performed similar to what described for Figure 6.

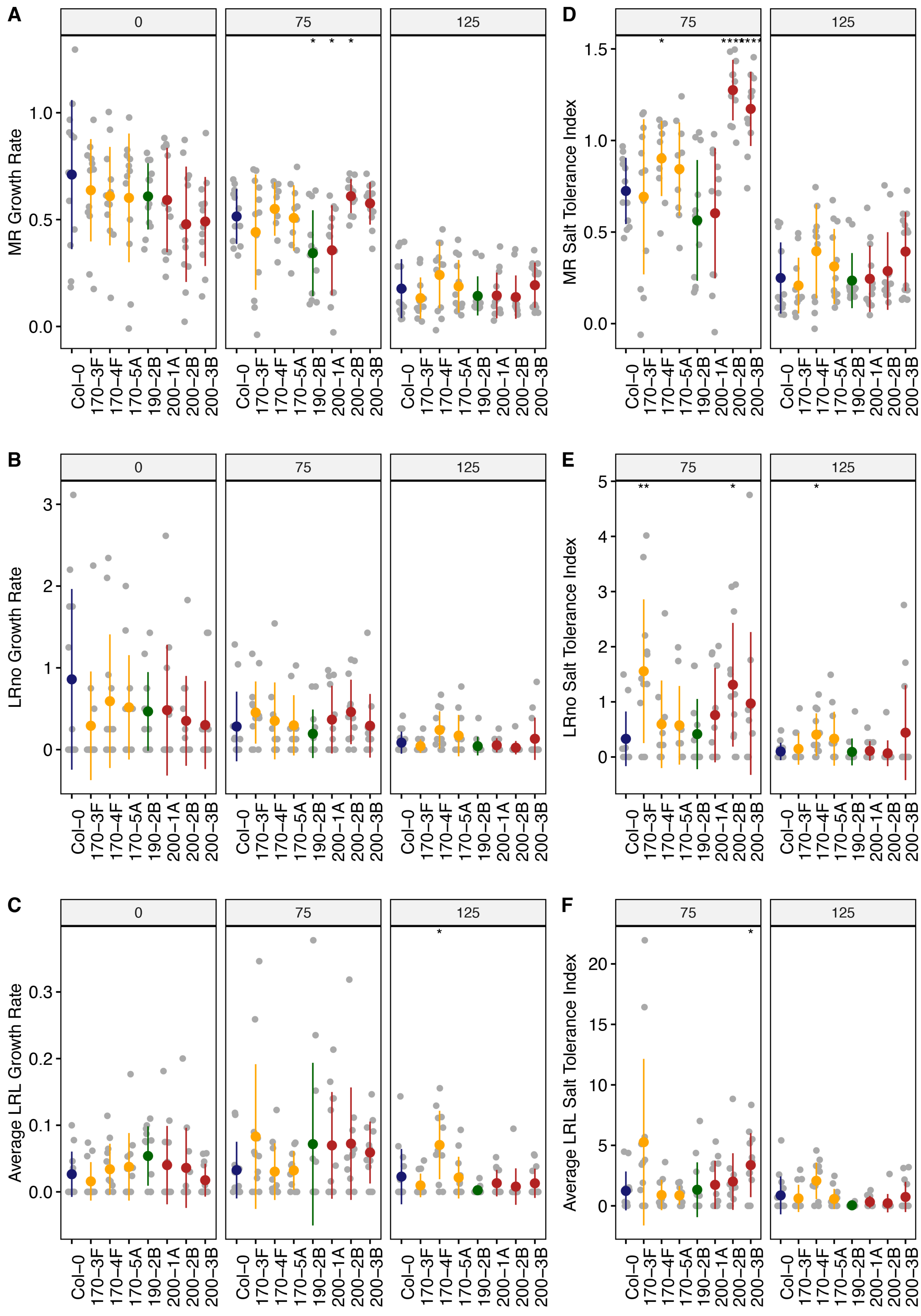


**Figure S20. Root system architecture analysis of neighboring DUF247 mutants under the salt, continues.** **(A)** Main root length, **(B)** lateral root number, and **(C)** lateral root length are shown for neighboring of the SR3G, including At3g50170 (shown as 170, with three T-DNA representatives, 170-3F (SALK_008710), 170-4F (SALK_145999), 170-5A (SALK_072937)), At3g50190 (shown as 190, with one T-DNA representative, 190-2B (SALK_137791)), and At3g50200 (shown as 200, with three T-DNA representatives, 200-1A (SALK_113759), 200-2B (SALK_093701), and 200-3B (SALK_129634)) compared to Col-0 under various concentrations of NaCl as indicated. The corresponding Salt Tolerance Index (STI) was calculated for **(D)** main root growth, **(E)** increase in lateral root number, and **(F)** lateral root growth rate. Statistical analysis was performed between Col-0 and the other mutants using the Student 's t-test: *P < 0.05. The root system architecture analysis was performed similar to what described for Figure 6.

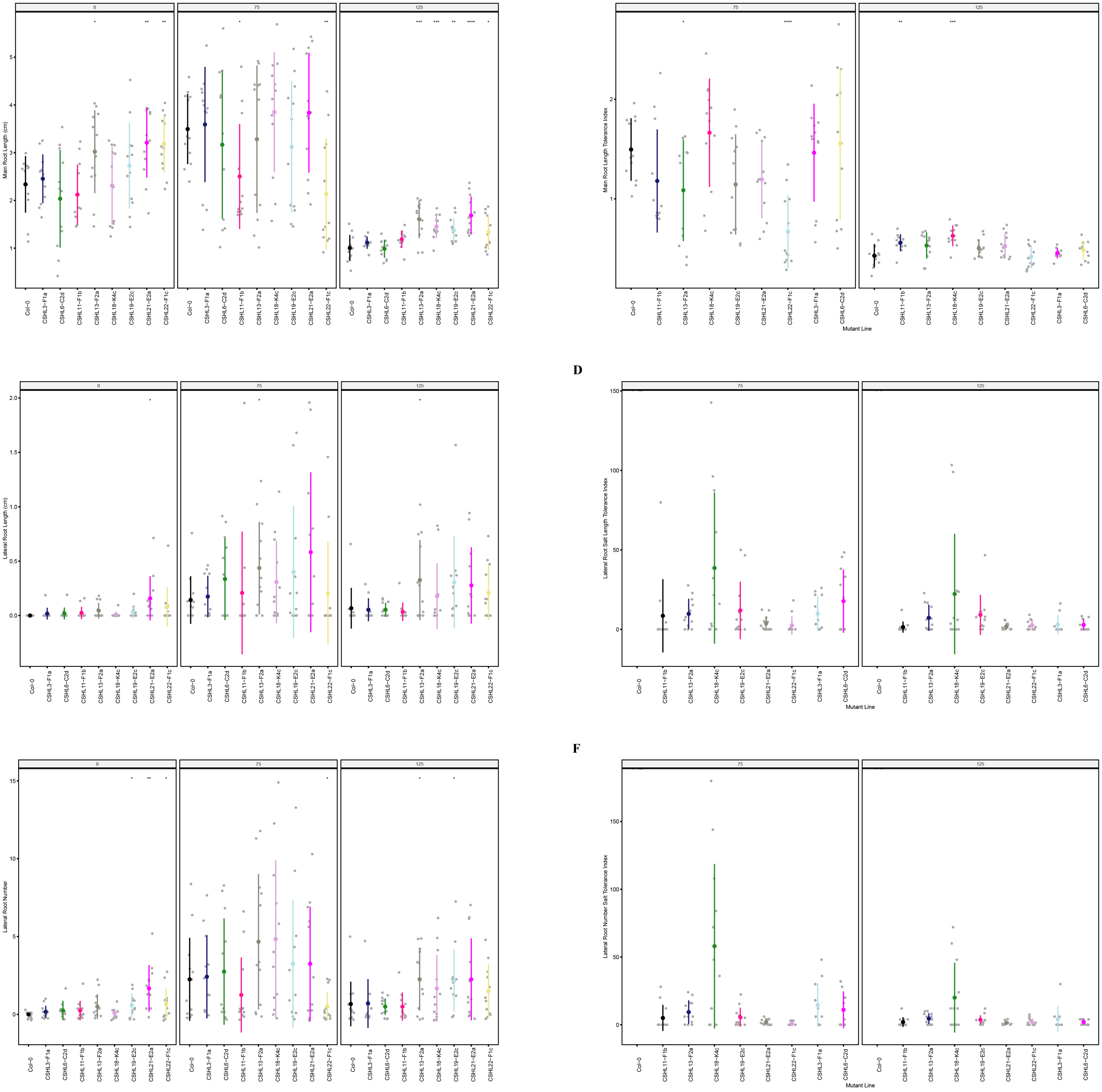


**Figure S21. Root system architecture analysis of RNAi lines targeting neighboring DUF247s. (A)** Main root length, **(B)** lateral root number, and **(C) l**ateral root length; **(D)** lateral root length tolerance index, **(E)** lateral root number, and **(F)** lateral root number tolerance index are shown for RNAi lines targeting AT3G50140 (CSHL3-F1a), targeting AT3G50130 and AT3G50140 (CSHL6-C2d), targeting AT3G50150 and AT3G50170 (CSHL11-F1b), targeting AT3G50140 and AT3G50170 (CSHL13-F2a), targetingAT3G50150 and AT3G50190 (CSHL18-K4c), targeting AT3G50130, AT3G50140, AT3G50150, AT3G50160, AT3G50170, AT3G50180, and AT3G50190 (CSHL19-E2c), targeting AT3G50130 and AT3G50150 (CSHL21-E2a), and targeting AT3G50150 and AT3G50160 (CSHL22-F1c). Statistical analysis was performed between Col-0 and the RNAi lines using a Student's t-test: *P < 0.05, **P < 0.01, ***P < 0.001, ****P < 0.0001.

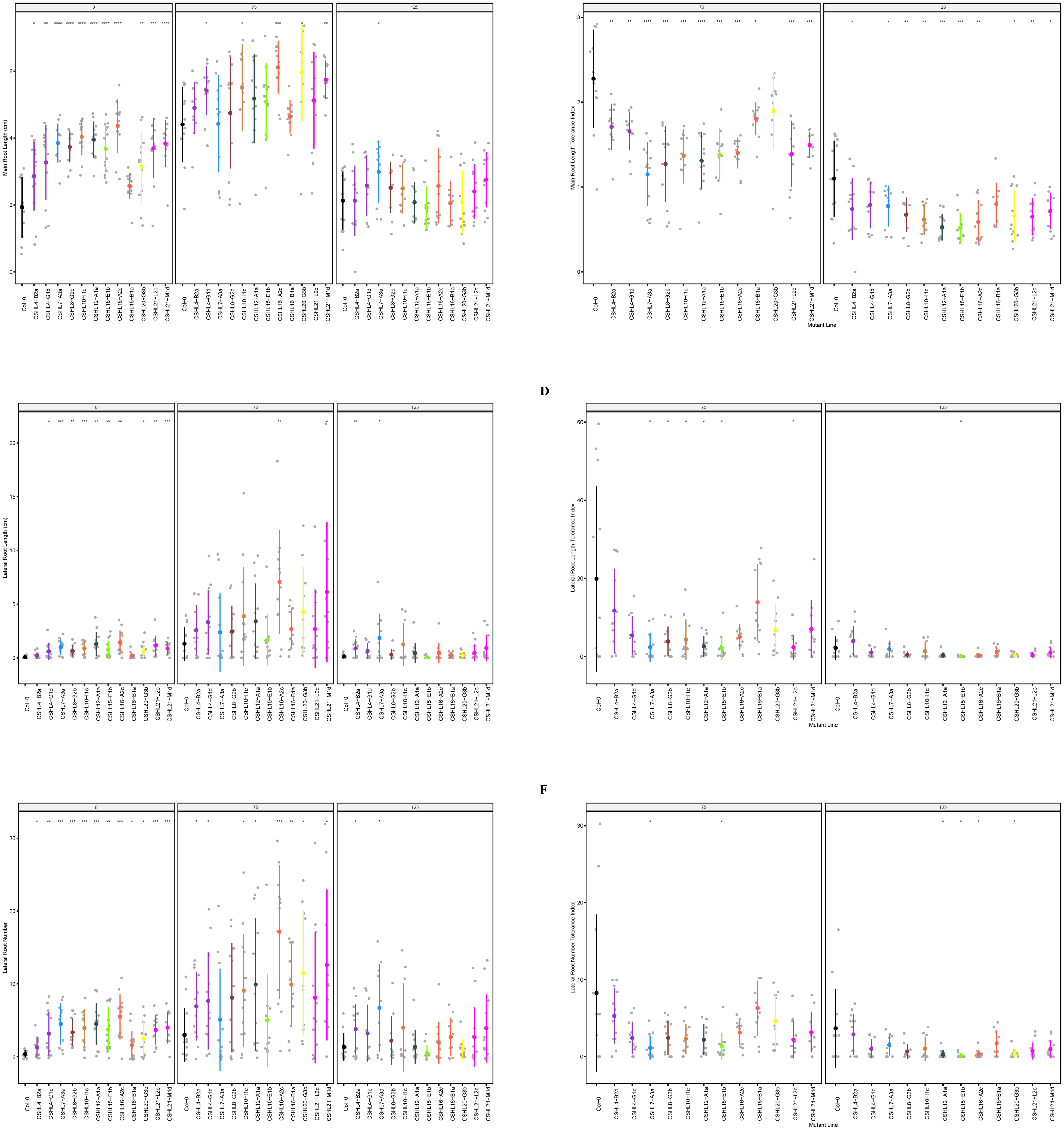


**Figure S22. Root system architecture analysis of RNAi lines targeting neighboring DUF247s continued. (A)** Main root length, **(B)** lateral root number, and **(C)** lateral root length; **(D)** lateral root length tolerance index, **(E)** lateral root number, and **(F)** lateral root number tolerance index are shown for RNAi lines targeting AT3G50160 (CSHL4-B2a and CSHL4G1d), targeting AT3G50140 and AT3G50150 (CSHL7-A3a), targeting AT3G50140 and AT3G50190 (CSHL8-G2b), targeting AT3G50180 (CSHL10-I1c), targeting AT3G50130, AT3G50140, AT3G50170, AT3G50180, and AT3G50190 (CSHL12-A1a), targeting AT3G50140, AT3G50170, AT3G50180, and AT3G50190 (CSHL15-E1b), targeting AT3G50130, AT3G50160, and AT3G50170 (CSHL-A2c and CSHL16-B1a), targeting AT3G50130 and AT3G50160 (CSHL20-G3b), and targeting AT3G50130 and AT3G50150 (CSHL21-L2c and CSHL21-M1d). Statistical analysis was performed between Col-0 and the RNAi lines using a Student's t-test: $*P < 0.05$, $**P < 0.01$, $***P < 0.001$, $****P < 0.0001$.

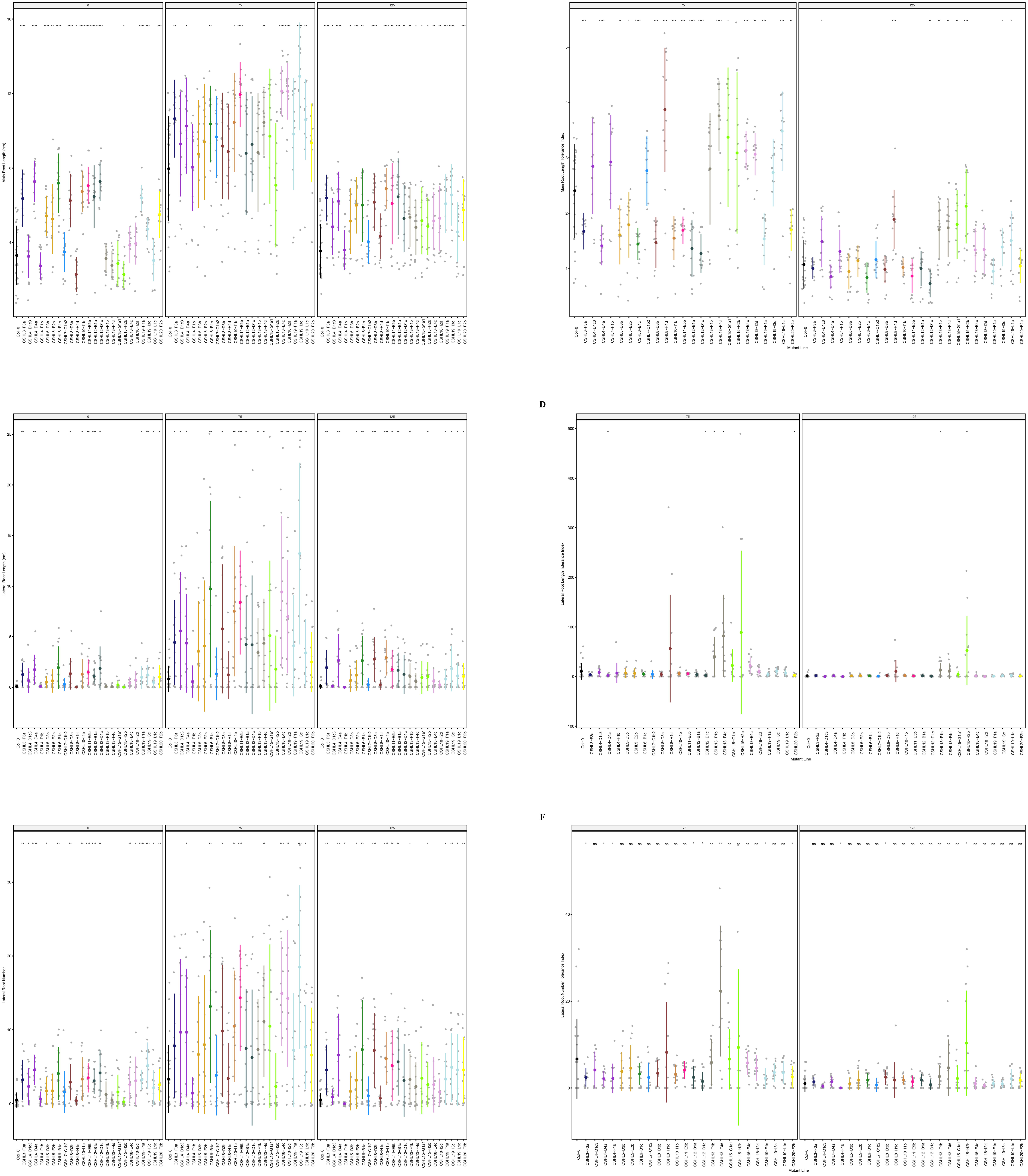


**Figure S23. Root system architecture analysis of RNAi lines targeting neighboring DUF247s continued. (A)** Main root length, **(B)** lateral root number, and **(C)** lateral root length; **(D)** lateral root length tolerance index, **(E)** lateral root number, and **(F)** lateral root number tolerance index are shown for RNAi lines targeting AT3G50140 (CSHL3-F3a), targeting AT3G50160 (CSHL4-D1c3, CSHL4-D4a, and CSHL4-F1b), targeting AT3G50130 and AT3G50170 (CSHL5-D3b and CSHL5-E2b), targeting AT3G50130 and AT3G50140 (CSHL6-B1c), targeting AT3G50140 and AT3G50150 (CSHL7-C1b2), targeting AT3G50140 and AT3G50190 (CSHL8-D3b and CSHL8-H1d), targeting AT3G50180 (CSHL10-I1b), targeting AT3G50150 and AT3G50170 (CSHL11-E5b), targeting AT3G50130, AT3G50140, AT3G50170, AT3G50180, and AT3G50190 (CSHL12-B1a and CSHL12-D1c), targeting AT3G50140 and AT3G50170 (CSHL13-F1b and CSHL13-F4d), targeting AT3G50140, AT3G50170, AT3G50180, and AT3G50190 (CSHL15-G1a1 and CSHL15-H2b), targeting AT3G50150 and AT3G50190 (CSHL18-E4c and CSHL18-I2d), targeting AT3G50130, AT3G50140, AT3G50150, AT3G50160, AT3G50170, AT3G50180, and AT3G50190 (CSHL19-F1a, CSHL19-I3c, and CSHL19-L1c), and targeting AT3G50130 and AT3G50160 (CSHL20-F2b). Statistical analysis was performed between Col-0 and the RNAi lines using a Student's t-test: *P < 0.05, **P < 0.01, ***P < 0.001, ****P < 0.0001.

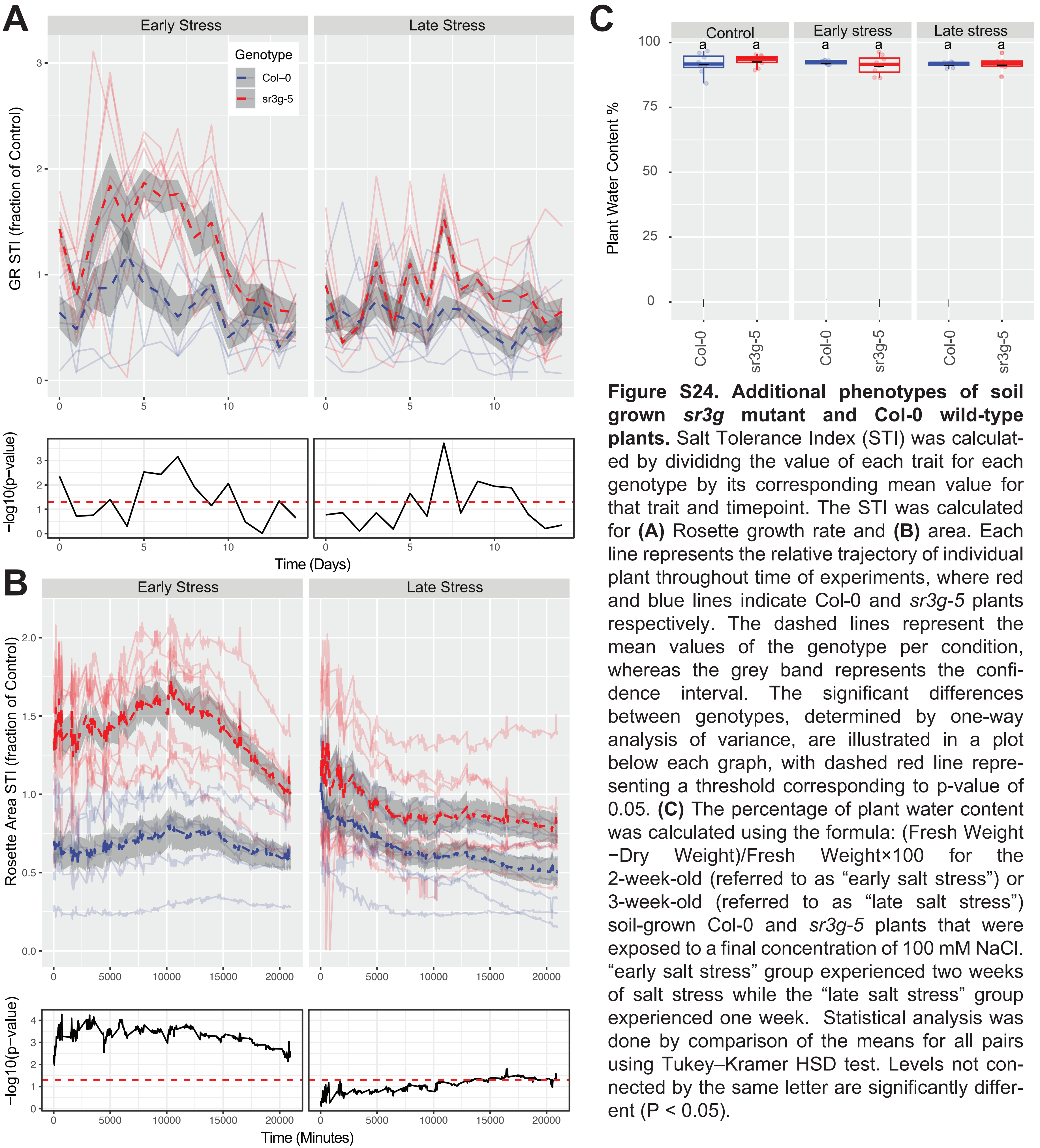


**Figure S24. Additional phenotypes of soil grown *sr3g* mutant and Col-0 wild-type plants.** Salt Tolerance Index (STI) was calculated by divididng the value of each trait for each genotype by its corresponding mean value for that trait and timepoint. The STI was calculated for **(A)** Rosette growth rate and **(B)** area. Each line represents the relative trajectory of individual plant throughout time of experiments, where red and blue lines indicate Col-0 and *sr3g-5* plants respectively. The dashed lines represent the mean values of the genotype per condition, whereas the grey band represents the confidence interval. The significant differences between genotypes, determined by one-way analysis of variance, are illustrated in a plot below each graph, with dashed red line representing a threshold corresponding to p-value of 0.05. **(C)** The percentage of plant water content was calculated using the formula: (Fresh Weight −Dry Weight)/Fresh Weight×100 for the 2-week-old (referred to as "early salt stress") or 3-week-old (referred to as "late salt stress") soil-grown Col-0 and *sr3g-5* plants that were exposed to a final concentration of 100 mM NaCl. "early salt stress" group experienced two weeks of salt stress while the "late salt stress" group experienced one week. Statistical analysis was done by comparison of the means for all pairs using Tukey–Kramer HSD test. Levels not connected by the same letter are significantly different ($P < 0.05$).

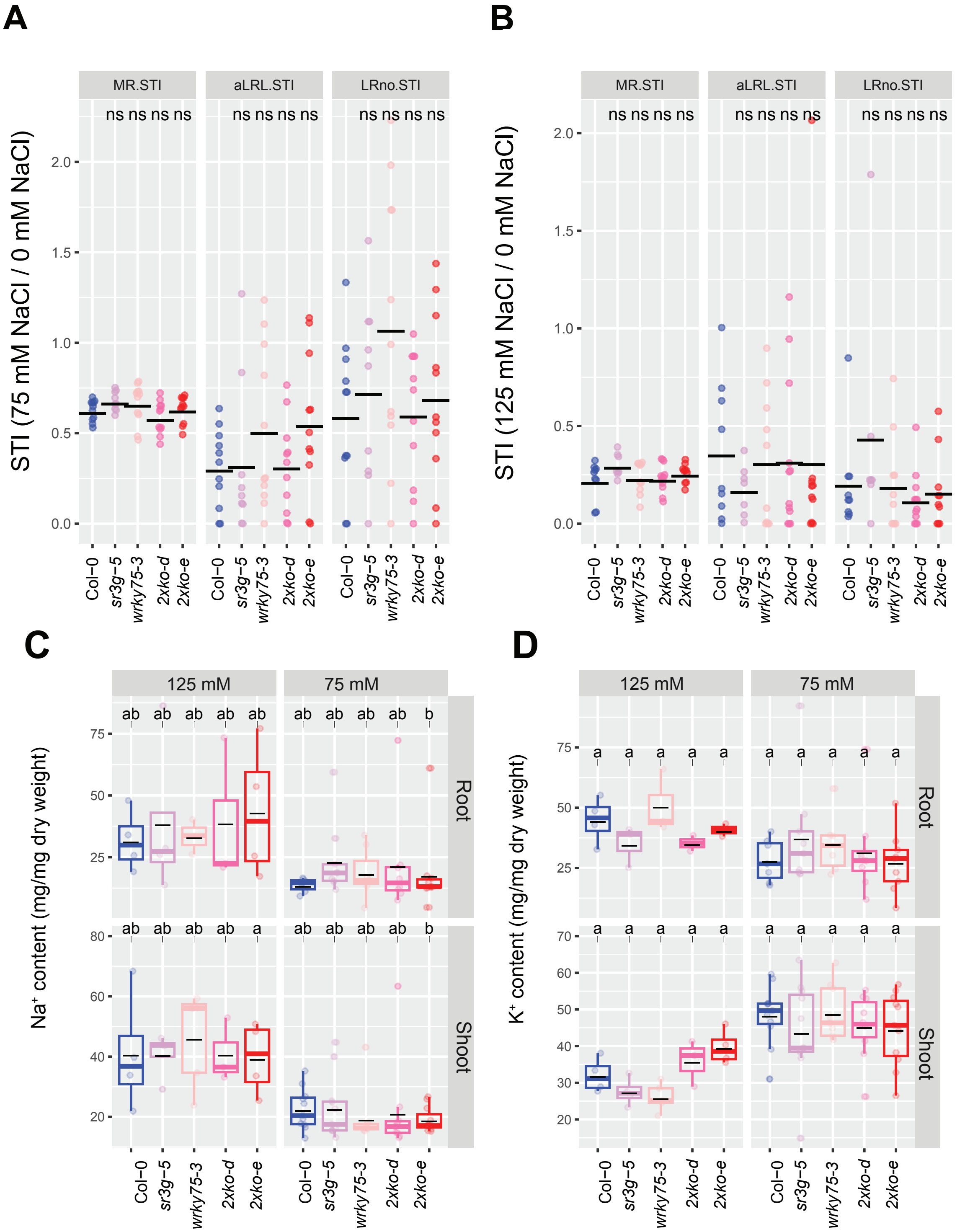


**Figure S25. wrky75/sr3g double mutants exhibit higher shoot K+ contents compared to Col-0 and each individual single mutant in the presence of 125 mM salt.** **(A)** Salt Tolerance Index (STI) for the main root length (MR.STI), average lateral root length (aLRL.STI), and lateral root number (LRno.STI) at 75 mM and **(B)** 125 mM NaCl are shown here. The STI was calculated by dividing the growth rate measured under salt stress by the growth rate measured under control condition for each genotype. The STI reported here were correspond to root system architecture analysis in Figure 10. Each dot in (A) and (B) represents individual replicate per genotype. In (A) and (B), significant differences between Col-0 and the other genotypes was determined by the Student 's t-test: *P < 0.05. ns stands for not significant. **(C)** $Na^+$ and **(D)** $K^+$ contents in root and shoot of Col-0, *sr3g-5*, *wrky75-3*, and two double mutants under various concentrations of NaCl are shown here. Statistical analysis in (A-B) was done by comparison of the means for all pairs using Tukey–Kramer HSD test. Levels not connected by the same letter are significantly different (P < 0.05).

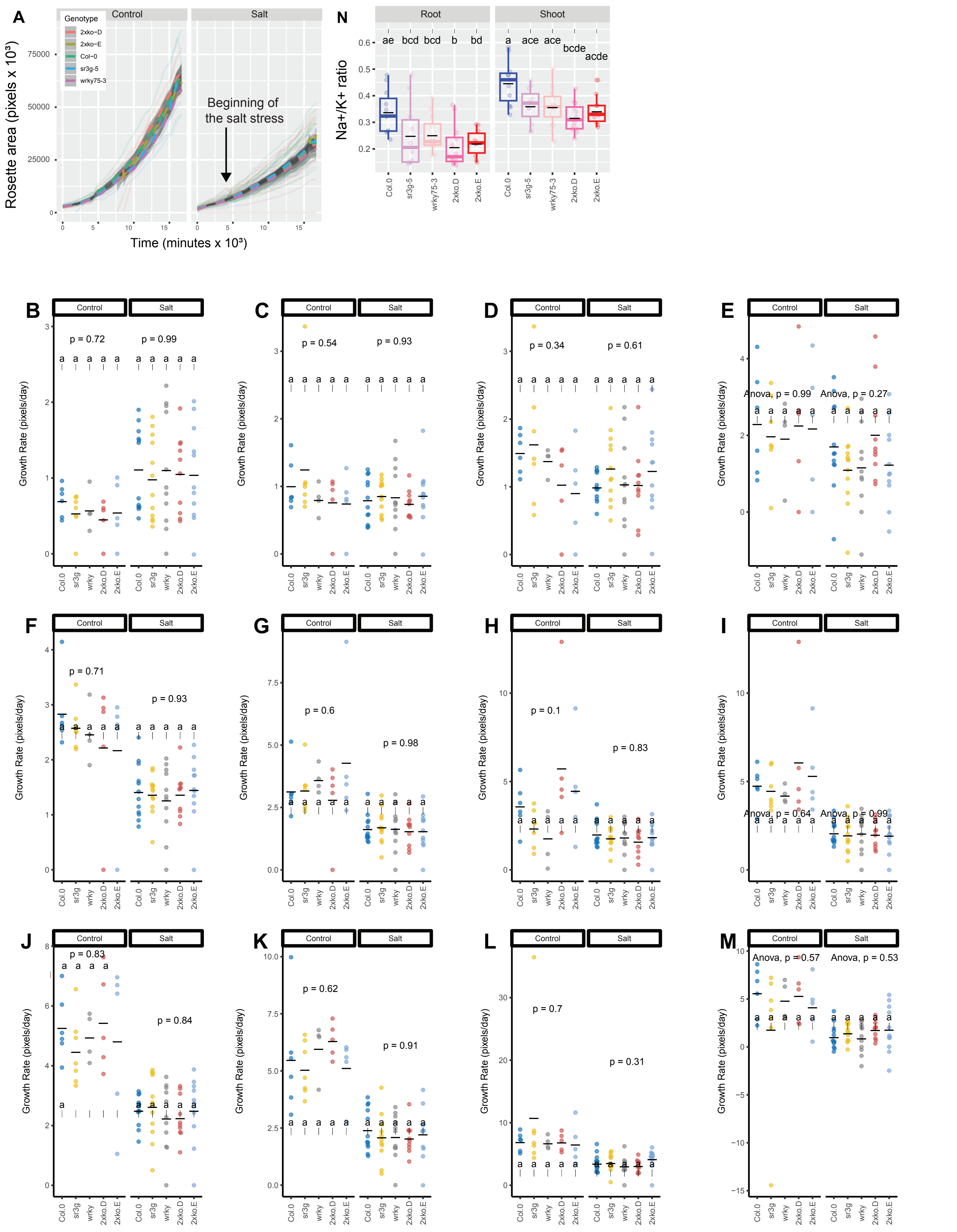


**Figure S26. wrky75/sr3g double mutants exhibit no noticeable differences in rosette area compared to their individual single mutants when grown in saline soil.** The salt stress responses of 2-week-old soil-grown plants that were exposed to a final concentration of 100 mM NaCl were examined here. **(A)** Rosette area and **(B-M)** growth rate for each individual day were shown . Growth rate is shown for **(B)** 1, **(C)** 2, and **(D)** 3 days before the stress, respectively. Growth rate is shown for **(E)** 1 , **(F)** 2, **(G)** 3 , **(H)** 4 , **(I)** 5, **(J)** 6, **(K)** 7, **(L)** 8, and **(M)** 9 days of stress application, respectively. **(N)** $Na^+/K^+$ ratio in shoot were measured in 4-week-old plants. Each line in (A) represents the trajectory per genotype through time. The effect of the genotype within individual treatment in (B-M) was tested using ANOVA, and the significantly different genotypes were additionally determined using Tukey HSD test (pvalue $< 0.05$). Statistical analysis in (N) was done by comparison of the means for all pairs using Tukey–Kramer HSD test. Levels not connected by the same letter are significantly different ($P < 0.05$).
